# Permeant Fluorescent Probes Visualize the Activation of SARM1 and Uncover an Anti-neurodegenerative Drug Candidate

**DOI:** 10.1101/2021.02.24.432704

**Authors:** Wan Hua Li, Ke Huang, Yang Cai, Qian Wen Wang, Wen Jie Zhu, Yun Nan Hou, Sujing Wang, Sheng Cao, Zhi Ying Zhao, Xu Jie Xie, Yang Du, Chi-Sing Lee, Hon Cheung Lee, Hongmin Zhang, Yong Juan Zhao

**Affiliations:** State Key Laboratory of Chemical Oncogenomics, Key Laboratory of Chemical Genomics, Peking University Shenzhen Graduate School, Shenzhen, China, 518055; Ciechanover Institute of Precision and Regenerative Medicine, School of Life and Health Sciences, The Chinese University of Hong Kong, Shenzhen, China, 518172; Department of Chemistry, Hong Kong Baptist University, Kowloon Tong, Hong Kong SAR, China; Department of Biology, and Shenzhen Key Laboratory of Cell Microenvironment, South University of Science and Technology of China, Shenzhen, China, 518055; Kobilka Institute of Innovative Drug Discovery, School of Life and Health Sciences, The Chinese University of Hong Kong, Shenzhen, China, 518172; Shenzhen-Hong Kong Institute of Brain Science-Shenzhen Fundamental Research Institutions, Shenzhen, China, 518055

## Abstract

SARM1 regulates axonal degeneration through its NAD-metabolizing activity and is a drug target for neurodegenerative disorders. We designed and synthesized fluorescent conjugates of styryl derivative with pyridine to serve as substrates of SARM1, which exhibited large red-shifts after conversion. With the conjugates, SARM1 activation was visualized in live cells following elevation of endogenous NMN or treatment with a cell-permeant NMN-analog. In neurons, imaging documented SARM1 activation preceded vincristine-induced axonal degeneration by hours. Library screening identified a derivative of nisoldipine as a covalent inhibitor of SARM1 that reacted with Cys311 in its Armadillo-domain and blocked its NMN-activation, protecting axons from degeneration. CryoEM showed that SARM1 was locked into an inactive conformation by the inhibitor, uncovering an unsuspected neuroprotective mechanism of dihydropyridines.

## Introduction

Axon degeneration (AxD) occurs in most neurodegenerative disorders (Coleman et al., 2020). Sterile Alpha and TIR Motif–containing 1 (SARM1) acts as a main effector in this process(Osterloh et al., 2012) and its depletion significantly attenuates AxD (Geisler et al., 2016, Osterloh et al., 2012, Turkiew et al., 2017). SARM1 controls AxD through its enzymatic activity(Essuman et al., 2017). It is self-inhibitory and is activated by nicotinamide mononucleotide (NMN)(Zhao et al., 2019), resulting in depletion of the intracellular NAD-pool(Essuman et al., 2017, Zhao et al., 2019). However, a recent study suggests that NAD may be an inhibitor of SARM1 activation and the decrease of NAD actually may activate SARM1 (Jiang et al., 2020). In any case, no direct evidence of SARM1 activation in live cells is available and no potent inhibitor has been identified. We thus aim to design and synthesize probes for visualizing SARM1-activation in live cells and to screen drug library for potent inhibitors.

We have documented that SARM1 is a multi-functional enzyme with properties similar to CD38, a universal signaling enzyme possessing not only NADase activity but also catalyzing both the cyclization of NAD to cyclic ADP-ribose (cADPR) and the exchange of nicotinamide in NADP with nicotinic acid to produce nicotinic acid adenine dinucleotide phosphate (NAADP) (Zhao et al., 2019). Both cADPR and NAADP are messengers regulating calcium mobilization in the endoplasmic reticulum and the endo-lysosomes, respectively (reviewed in(Galione, 1994, Lee, 2012, Lee et al., 2019)). The catalytic similarities and its ubiquitous presence in non-neuronal cells suggest that SARM1 may be a calcium signaling enzyme as well.

We focused on its base-exchange reaction for designing specific probes for SARM1 and had shown that pyridyl derivatives can readily serve as substrates (Graeff et al., 2006, Lee et al., 1997). We thus conjugated various styryl derivatives to pyridine to produce a series of conjugates (PCs) as fluorescent probes for SARM1 activity (Fig. 1*A*).

**Fig. 1.**
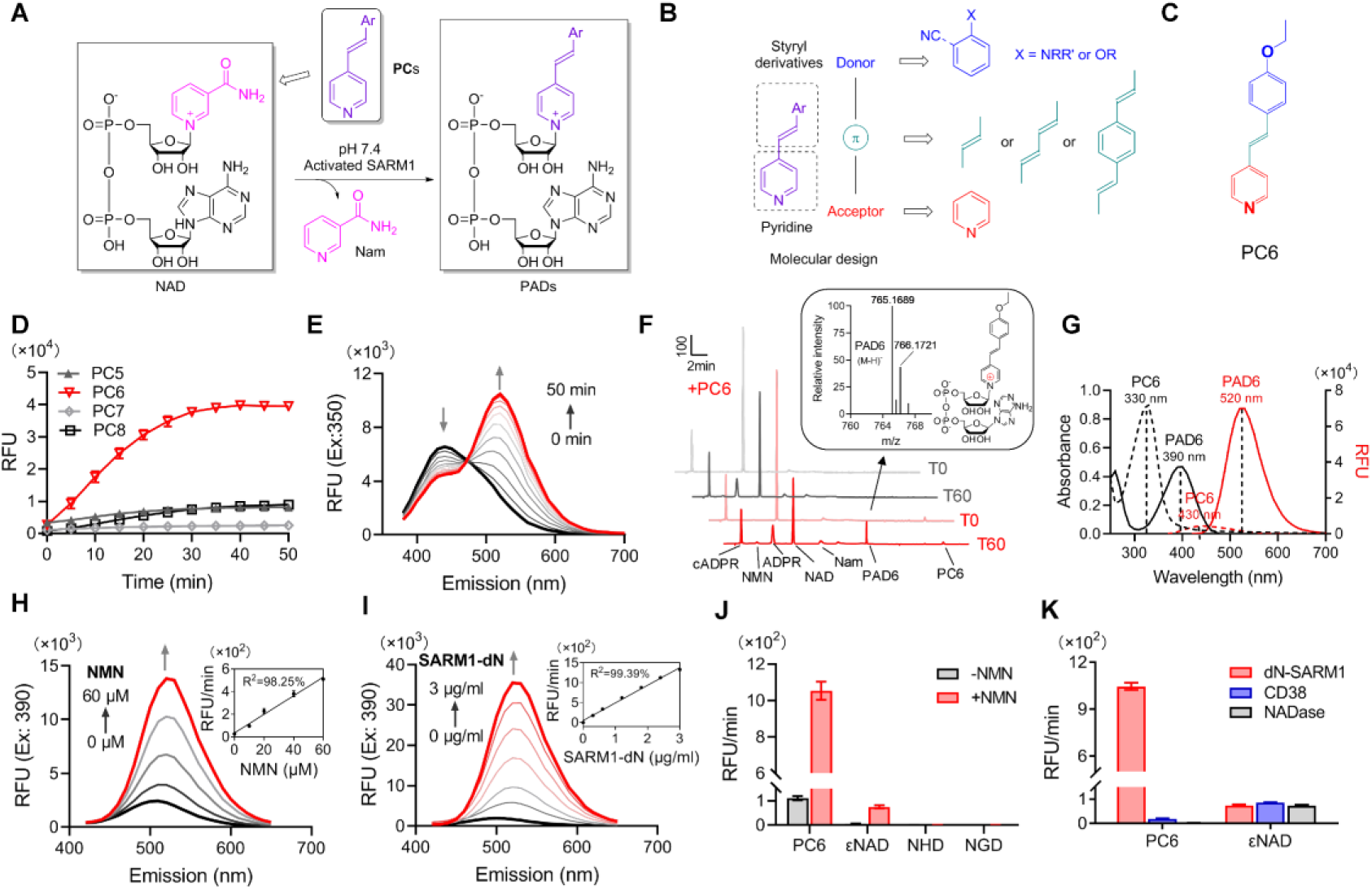
Design and characterization of PC probes. (*A*) Strategy of fluorescent imaging of the activated SARM1. (*B*) Designing based on pyridine and styryl derivatives with a donor-π-acceptor framework. (*C*) Structure of PC6. (*D*) The kinetics of the fluorescence increase at the maximal absorbance wavelengths catalyzed by SARM1-dN, in the presence of 100 μM NMN, 100 μM NAD and 50 μM PCs. (*E*) Time-dependent changes of the emission spectra at the isosbestic point (350 nm). (*F*) HPLC of PC6 reaction. Red line: in the presence of PC6, NMN and NAD; Gray line: without PC6. Insert: MS analysis and structure of PAD6. (*G*) Absorbance and fluorescence spectra of 25 μM PC6/PAD6. (*H*) Emission spectra with dose of NMN in the presence of NAD, PC6 and SARM1-dN. Inset: the initial rates plotted to NMN concentrations. (*I*) Emission spectra with dose of SARM1-dN in the presence of NMN, NAD and PC6; Inset: the initial rate plotted to SARM1 concentration. (*J*) The reaction rates of 10 μM PC6 compared with NAD analogues (100 μM) catalyzed by SARM1. (*K*) The reaction rates of 10 μM PC6 catalyzed by SARM1, NADase and CD38.

## Results

### Probe design, synthesis, and characterization

We reasoned that conjugating the electron-rich styryl derivative with pyridine should provide a donor-π-acceptor framework(Pawlicki et al., 2009) (Fig. 1*B*). The positive charge of the pyridinium moiety of the product should delocalize over the conjugated π-system and lead to fluorescence changes (Fig. 1*A*). Pyridine conjugates (PC1-9, Figure 1—figure supplement 1, Figure 1—figure supplement 2, Figure 1—figure supplement 3*A*) were synthesized using the Pd-catalyzed cross-coupling strategy with yields ranging from 33.5-85.0%. The synthesis details are in the Methods section and product characterizations are Figure 1—figure supplement 2.

The PC-probes were tested using a recombinant SARM1, SARM1-dN(Zhao et al., 2019) (described in Figure 1—figure supplement 3*B*), with NAD as the acceptor of base-exchange and NMN as an activator. As shown in Figure 1—figure supplement 4, significant shifts in UV-vis spectra after conversion were observed in the oxygenated derivatives (PC5-9, *O*-series), but not the nitrogenated derivatives (PC1-4, *N*-series). The emission spectra of the reactive *O*-series showed steady increase as the reaction progressed (Figure 1—figure supplement 5, spectra; Fig. 1*D*, kinetics; Figure 1—figure supplement 3*C*, initial rate), with PC6, the chemical structure in Fig. 1*C*, exhibited the largest fluorescence increase (Fig. 1*D*).

The time course of the UV-spectra during conversion of PC6 showed decreases at 330 nm but increases at 400, with an isosbestic point at 350 nm (Figure 1—figure supplement 4 and Fig. 1*E*). Corresponding to the absorbance change was the red-shift in the fluorescence spectra, from the emission maximum at 430 nm of PC6 to 520 nm of PAD6 (Fig. 1*E*).

The conversion of PC6 to the exchange-product, PAD6, was verified by purifying it using HPLC and characterized by high resolution mass spectrometry (HRMS) (Fig. 1*F*). The remarkably large spectral changes are anticipated from our design, as the pyridine ring becomes positively charged after its exchange into NAD (Fig. 1*F*, inlet), a much stronger electron acceptor in the D−π−A structure, thereby increasing intramolecular charge transfer and shifting the emission maximum by over 100 nm. The conversion-induced spectral changes were consistent with the spectra of the HPLC-purified products, PAD6 (Fig. 1*G*).

The observed spectral changes showed a linear dependence on NMN, with as low as 10 μM being effective (Fig. 1*H*), confirming that SARM1 is an auto-inhibitory enzyme activated by NMN(Zhao et al., 2019). The fluorescence increase was also proportional to the amount of NMN-activated SARM1 (Fig. 1*I*), with a detection limit of 48 ng/ml. As an *in vitro* assay for SARM1, PC6 provides more than 100-fold higher sensitivity over other commonly used probes, such as εNAD, NGD or NHD (Fig. 1*J*).

In addition to sensitivity, PC6 also shows exquisite selectivity toward SARM1 versus CD38 and *N. crassa* NADase(Graeff et al., 1994). All three possess NADase activity as detected by εNAD (Fig. 1*K*), but only SARM1 could produce large fluorescence increases with PC6.

The online version of this article includes the following figure supplements for figure 1:

**figure supplement 1.**
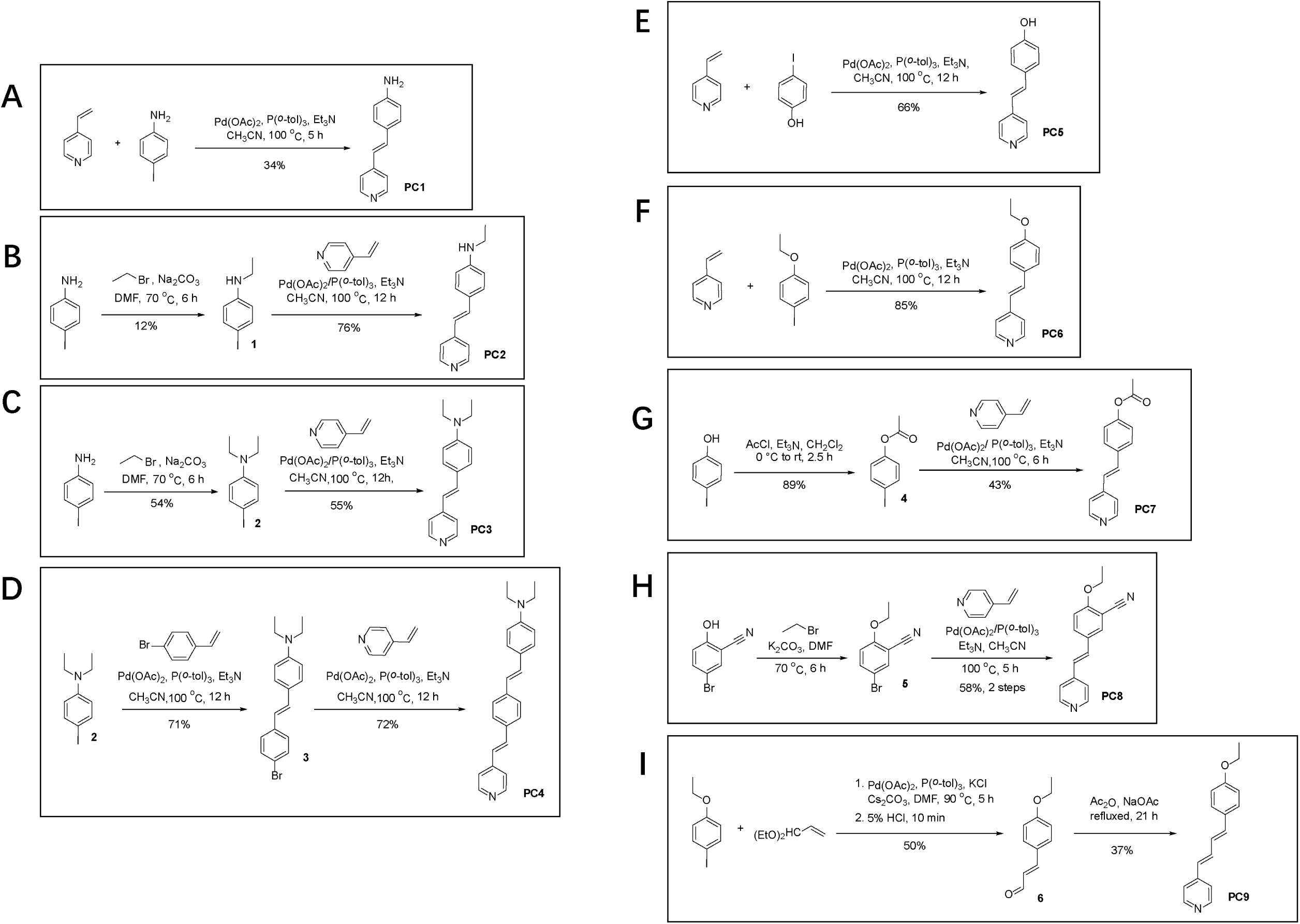
Synthetic scheme of PC1-PC11

**figure supplement 2.**
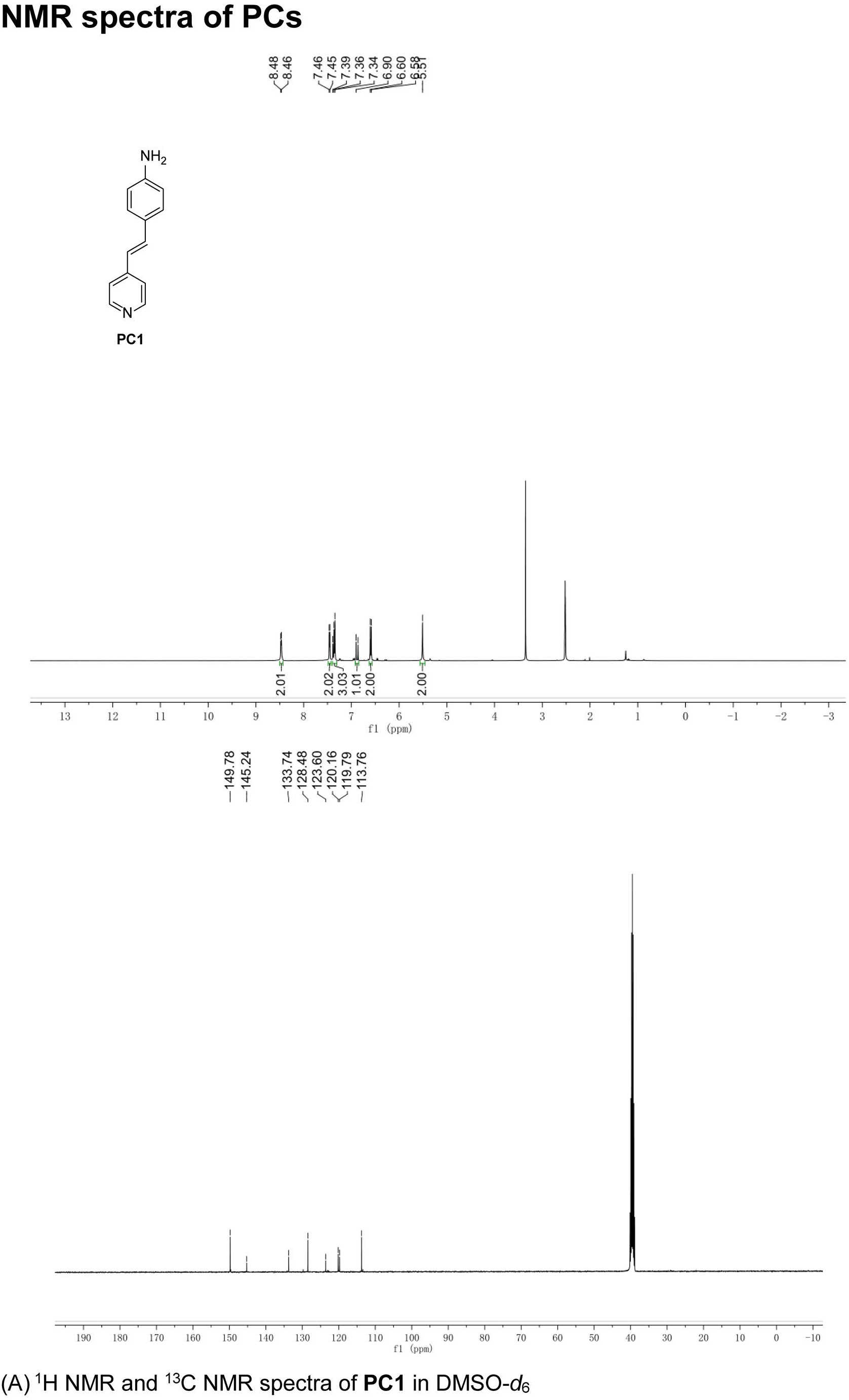
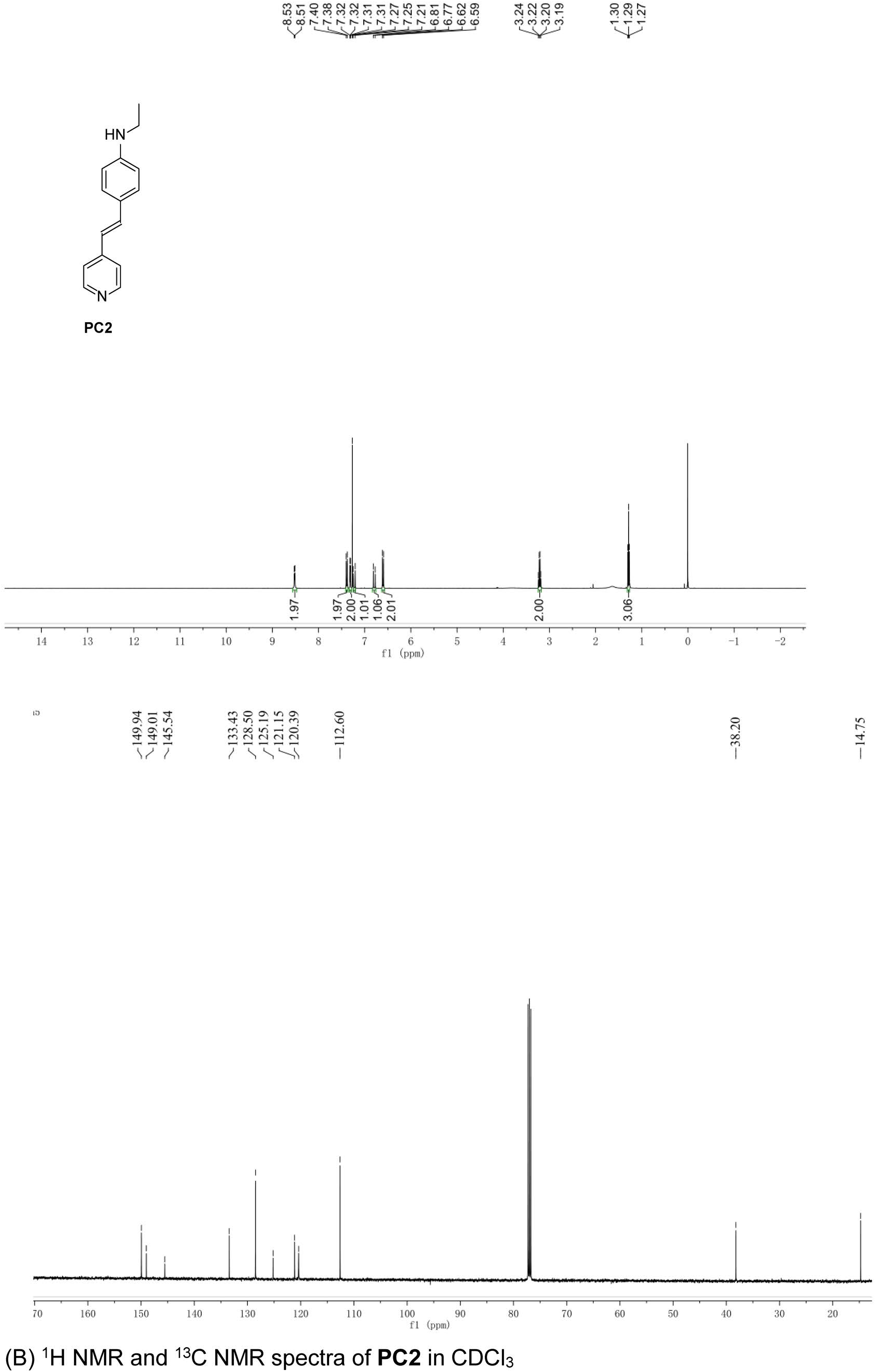
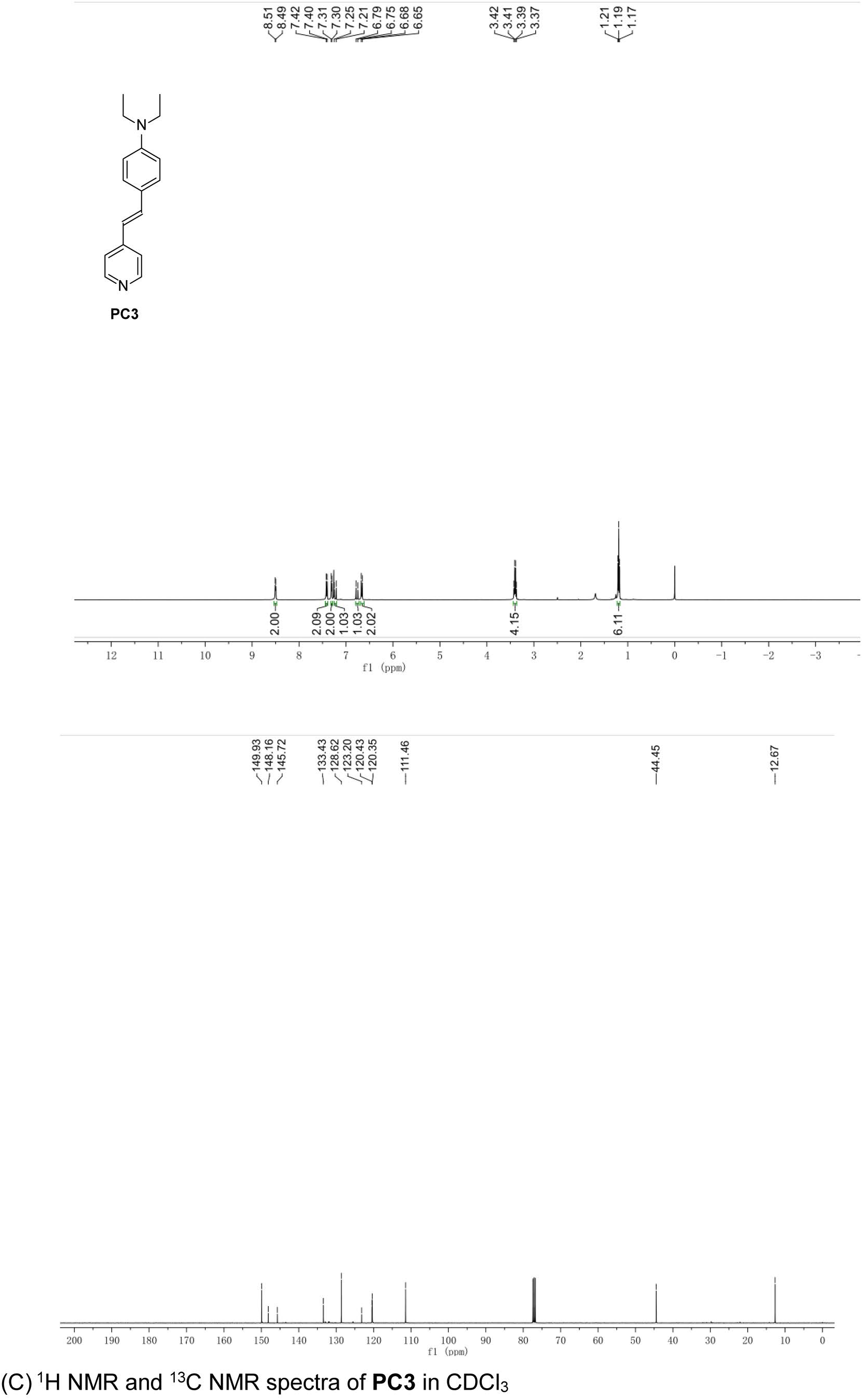
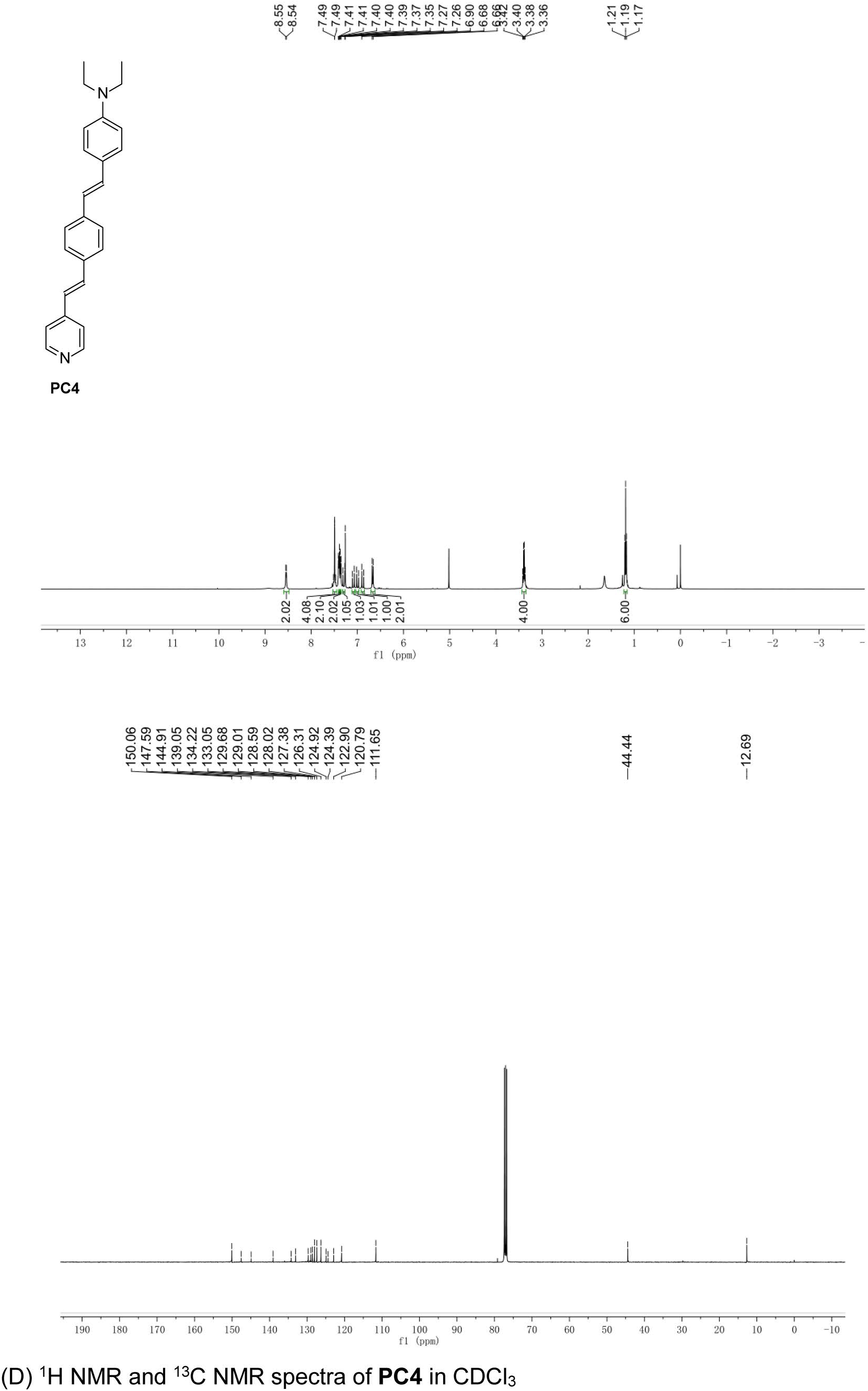
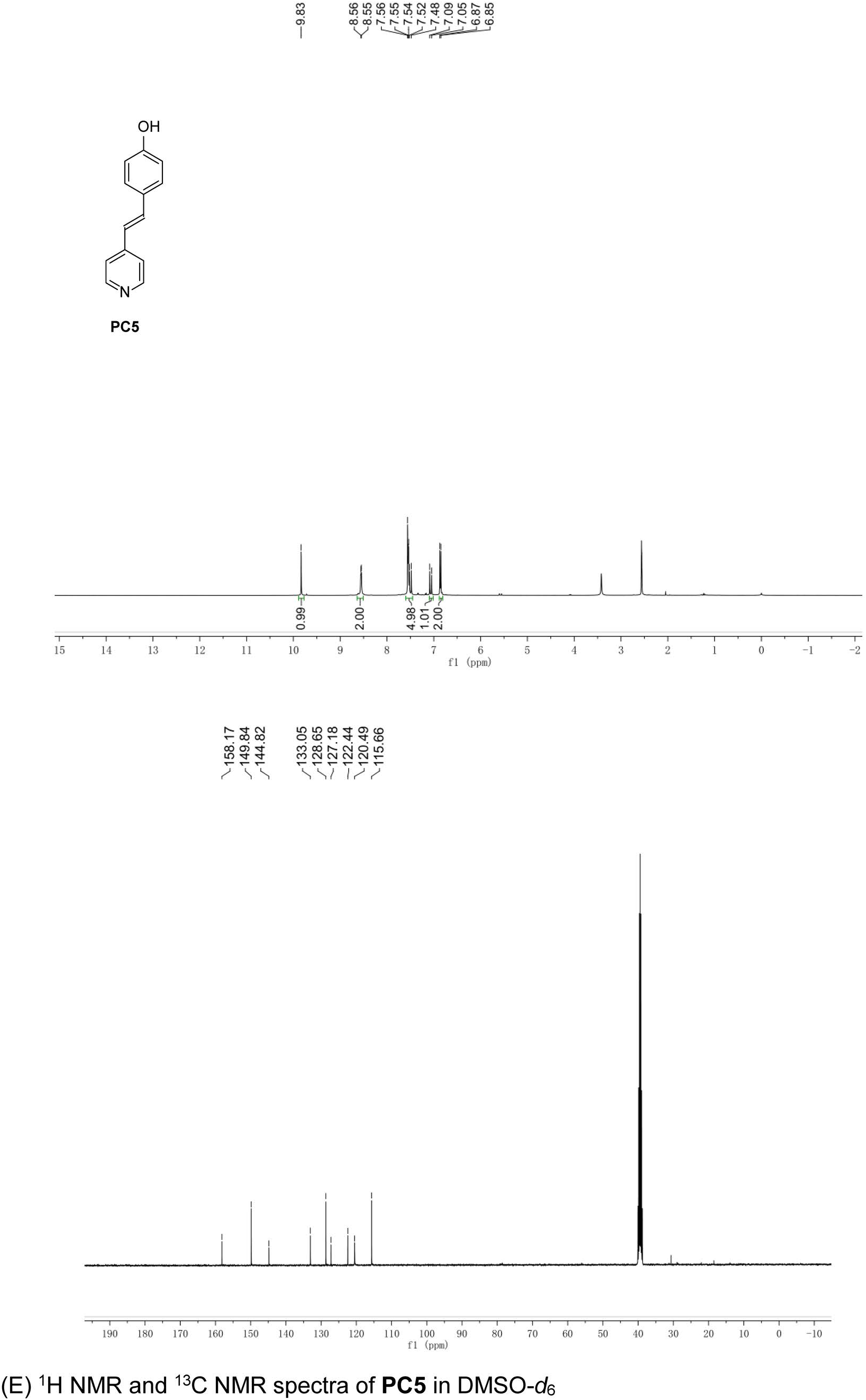
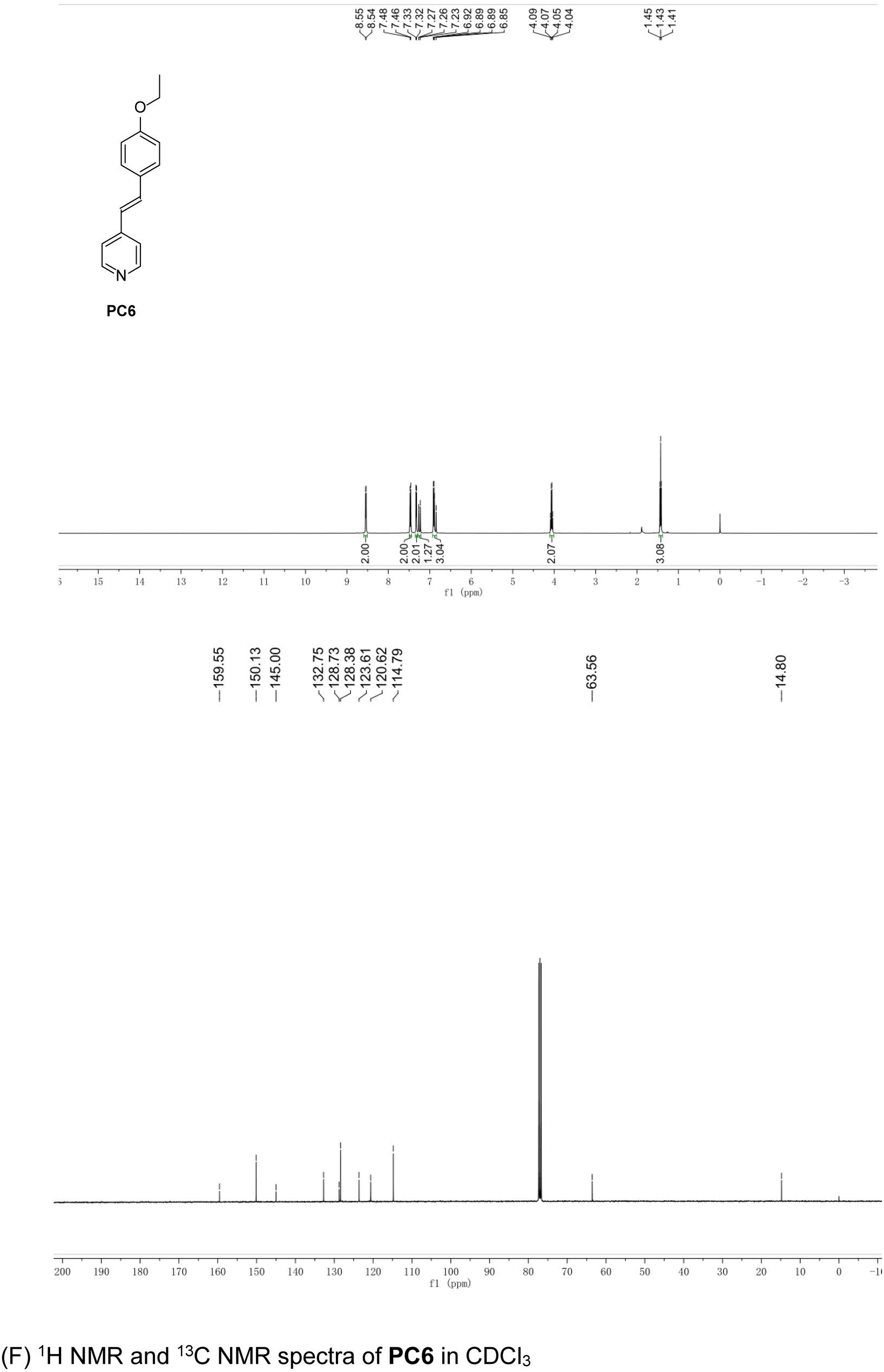
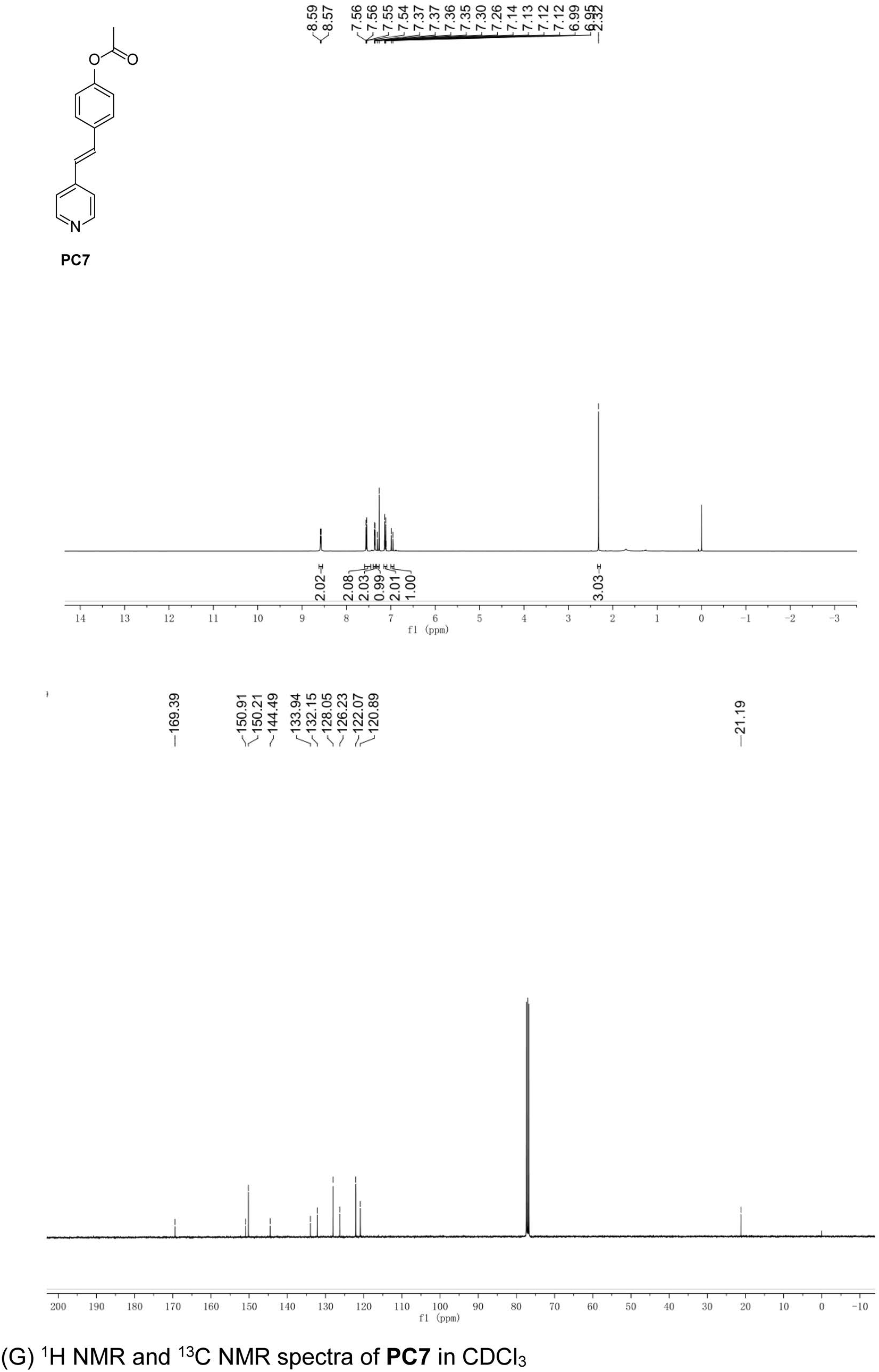
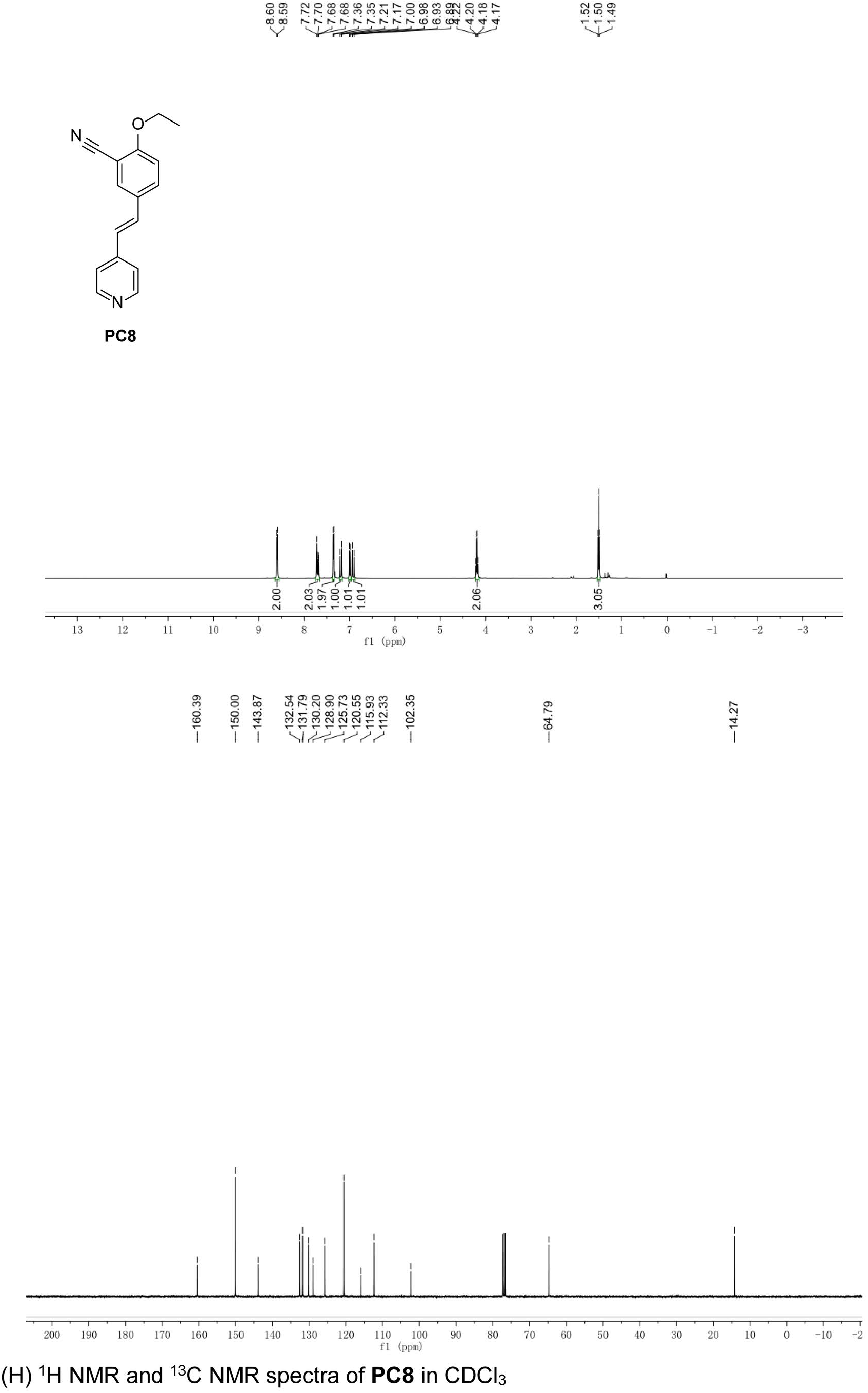
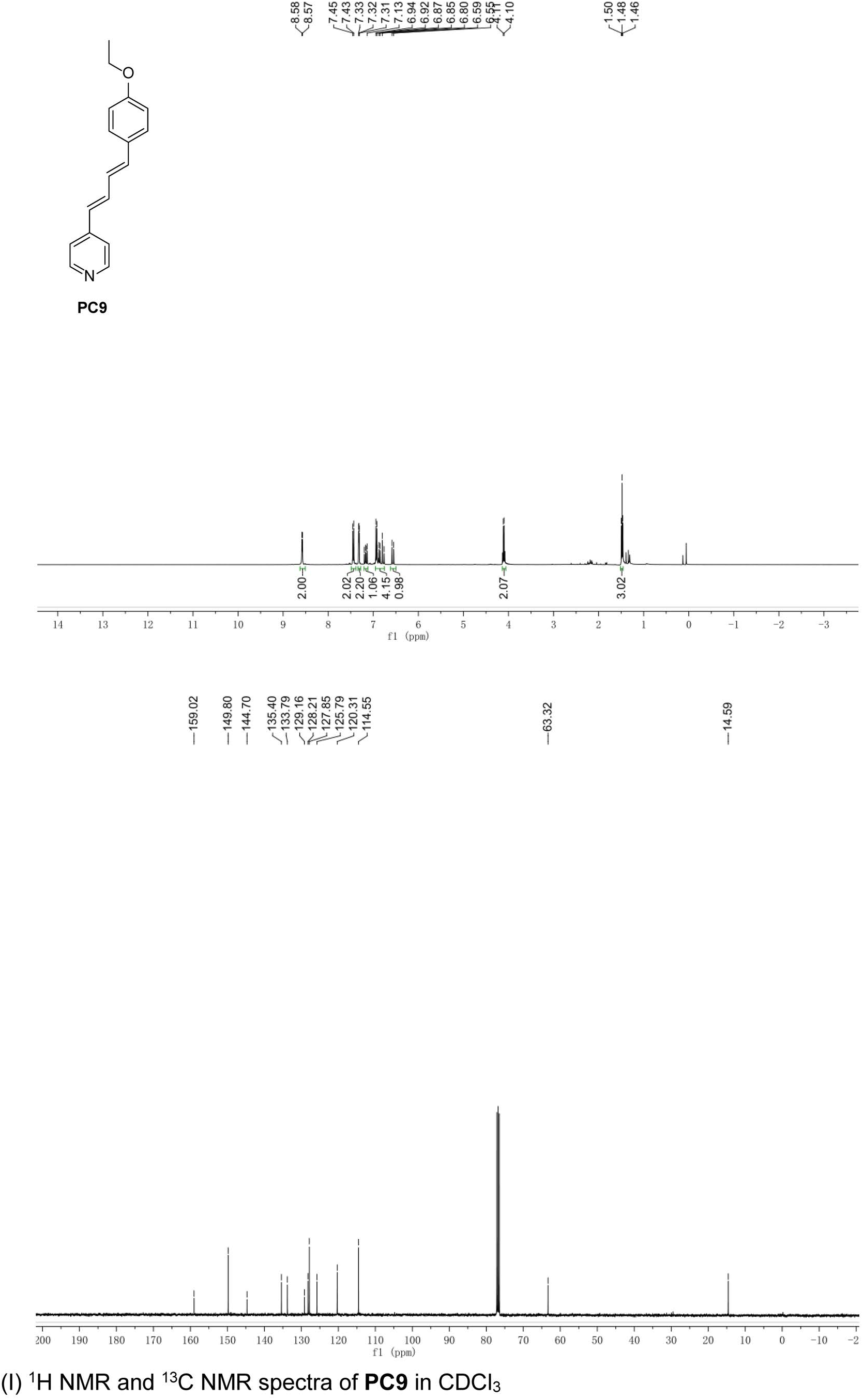
NMR spectra of PCs (A) ^1^H NMR and ^13^C NMR spectra of **PC1** in DMSO-*d*_6_ (B) ^1^H NMR and ^13^C NMR spectra of **PC2** in CDCl_3_ (C) ^1^H NMR and ^13^C NMR spectra of **PC3** in CDCl_3_ (D) ^1^H NMR and ^13^C NMR spectra of **PC4** in CDCl_3_ (E) ^1^H NMR and ^13^C NMR spectra of **PC5** in DMSO-*d*_6_ (F) ^1^H NMR and ^13^C NMR spectra of **PC6** in CDCl_3_ (G) ^1^H NMR and ^13^C NMR spectra of **PC7** in CDCl_3_ (H) ^1^H NMR and ^13^C NMR spectra of **PC8** in CDCl_3_ (I) ^1^H NMR and ^13^C NMR spectra of **PC9** in CDCl_3_

**Figure supplement 3.**
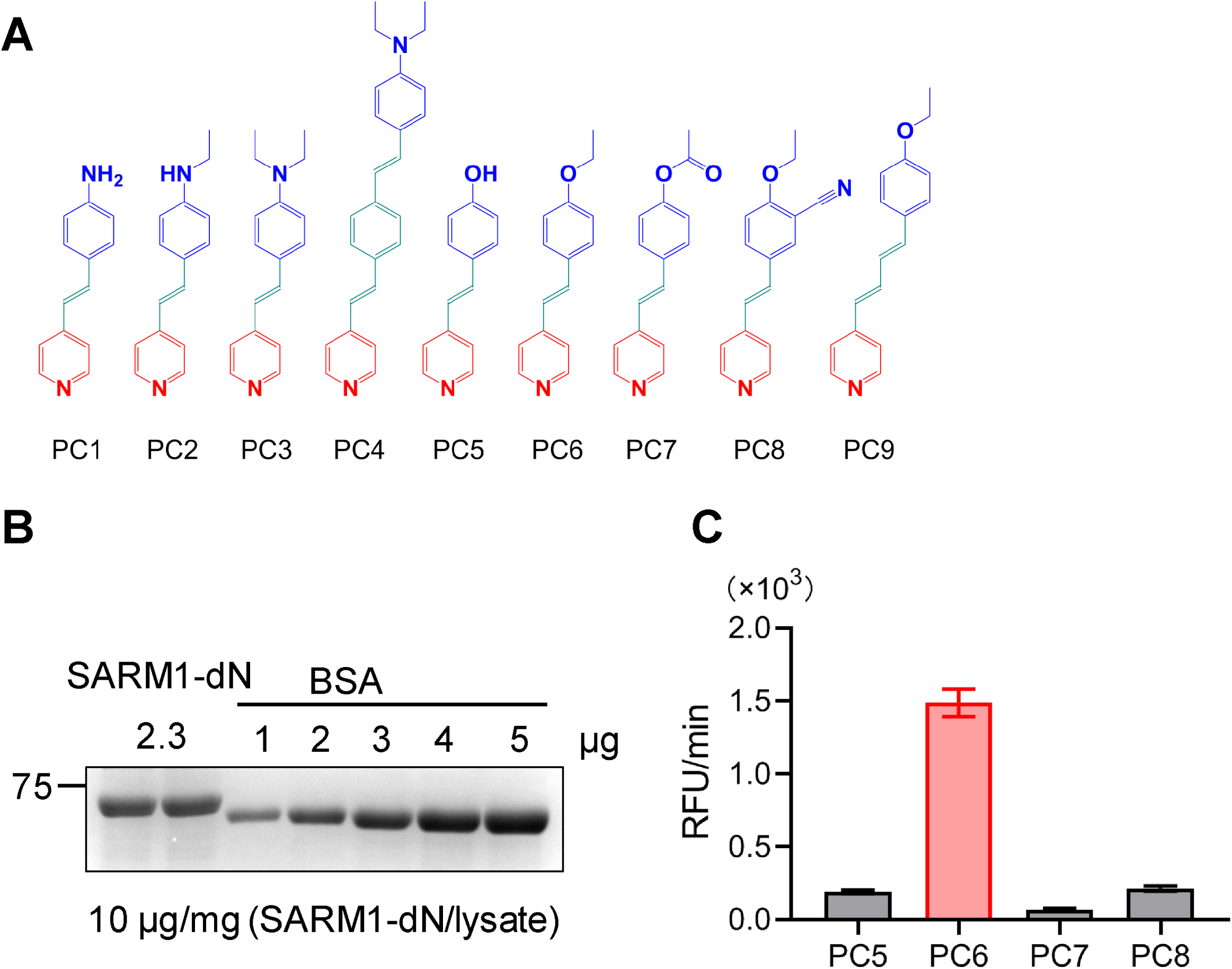
Structures of PC1-9 and activity screening. (*A*) Chemical structures of PC1-9. (*B*) quantification of SARM1-dN. SARM1-dN was pulled down by the BC2 nanobody(Zhao et al., 2019) conjugated beads, which efficiency was close to 100%. The purified SARM1-dN proteins were supplied to SDS-PAGE and Coomassie blue staining. The protein contents of SARM1-dN were calculated with a standard curve of BSA after intensity scanning. (*C*) The initial rates of the fluorescence increase at the maximal absorbance/emission wavelengths (PC5: 400nm/530nm; PC6: 390 nm/520 nm; PC7: 340 nm/445 nm; PC8: 375 nm/490 nm) catalyzed by SARM1-dN, in the presence of NMN, NAD and PCs.

**Figure supplement 4.**
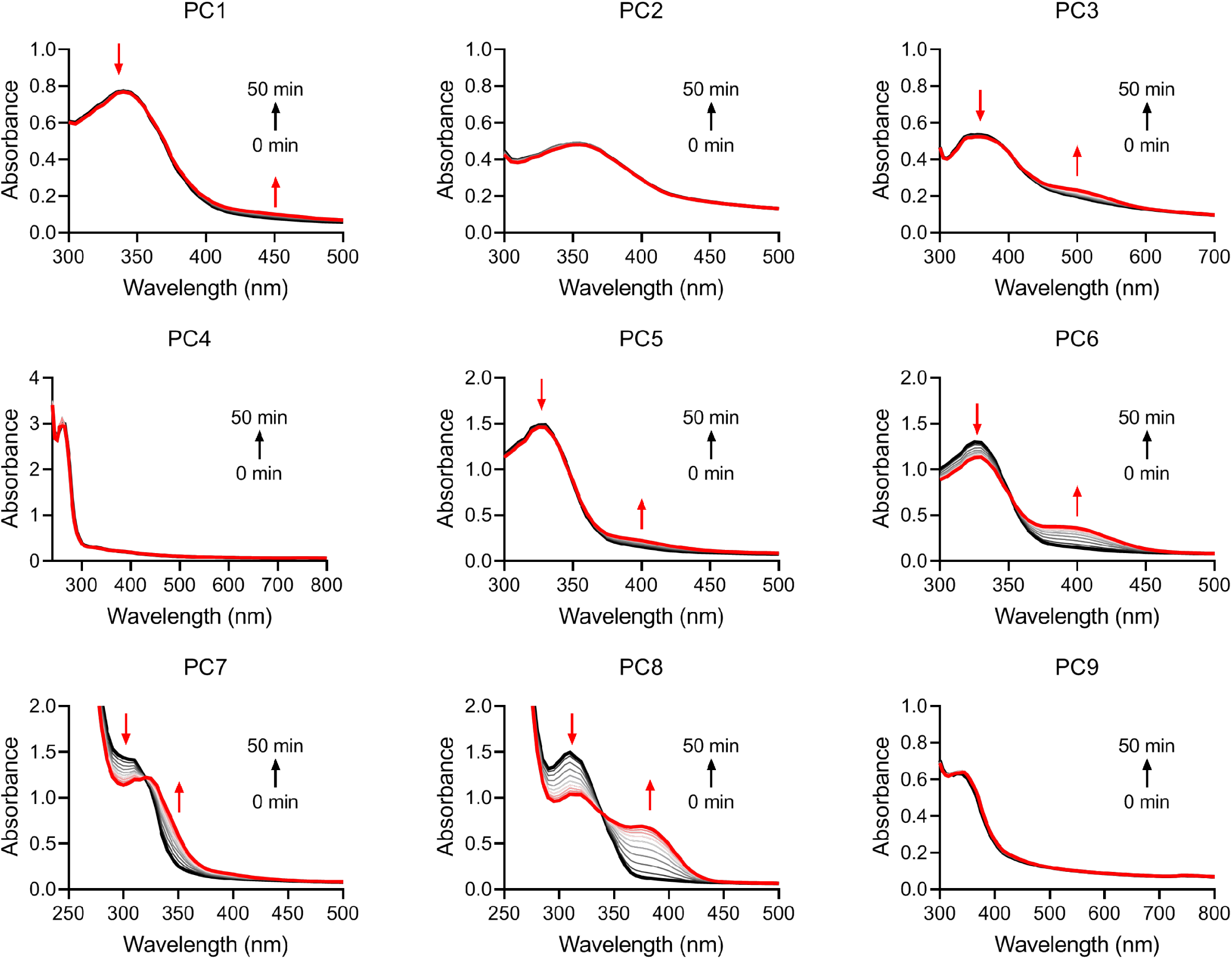
UV-vis absorption spectra scanning of the reactants. The reactants of 50 μM PCs and 100 μM NAD during the 50-min reactions catalyzed by the activated SARM1.

**Figure supplement 5.**
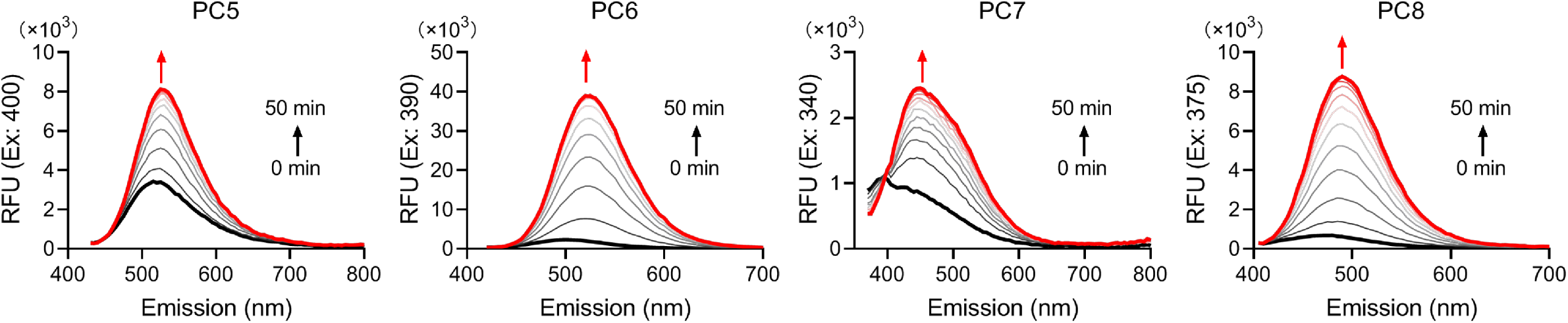
Fluorescence spectra of the reactants. The reactants of 50 μM PCs and 100 μM NAD during the 50-min reactions catalyzed by the activated SARM1.

### Imaging SARM1 activation in live cells

PC6 was added to HEK293 cells overexpressing either wildtype SARM1 or the enzymatically inactive mutant, E642A(Essuman et al., 2017, Zhao et al., 2019) (Fig. 2*A*). Green fluorescence was clearly seen evenly distributed in the whole cells in the wildtype, but not in the mutant cells (Fig. 2*B*), indicating active SARM1 was required. Intracellular production of PAD6 was confirmed in extracts of wildtype but not the E642A cells (Fig. 2*C*). CZ-48, a cell-permeant mimetic of NMN and activator of SARM1(Zhao et al., 2019), dramatically increased the PAD6 fluorescence (Fig. 2*B*, right column) and none in E642A-cells. These results indicate that PC6 is cell permeant and can be exchanged into the cytosolic NAD by the activated SARM1 to produce PAD6 having a large red-shift in fluorescence. PAD6 was also cell impermeant because of its charged ADP-ribose moiety and accumulated in the cytosol, greatly increased its detection sensitivity in live cells.

**Fig. 2.**
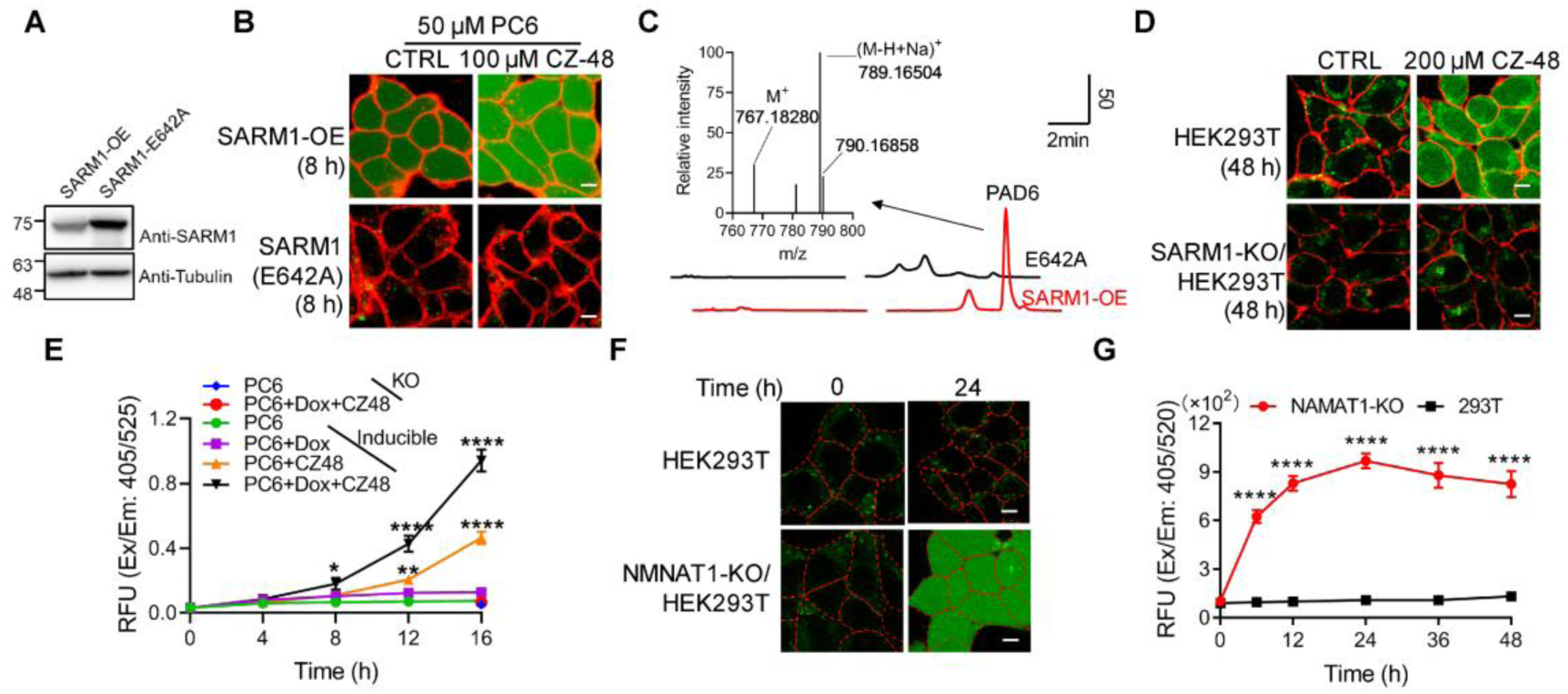
Live-cell imaging of SARM1 activation. (*A*) Western blot of the overexpression of SARM1 and inactive mutant, E642A in HEK293 cells. (*B*) Confocal fluorescence images of cells in (A) after incubation with PC6 in presence or absence of CZ-48. Green: PAD4; Red: ConA-Alex-647; (*C*) HPLC and MS analysis of PAD6 from SARM1-OE cells. The metabolites were extracted by 0.6 M PCA from the cells in (A) after treating with 50 μM PC6 for 24 h. Inset: MS analysis. (*D*) Confocal images of wildtype, or SARM1-KO HEK293T cells with PC6 in presence or absence of CZ-48. (*E*) The HEK293 cells carrying the inducible SARM1 were incubated with 50 μM PC6 in presence or absence of 0.5 mg/mL Dox and/or 100 μM CZ-48. The PAD6 fluorescence was analyzed by flow cytometry. (*F*) Confocal images of NMNAT1-KO/ HEK293T cells, incubated with PC6. Cell edges were marked according to the bright-field images. (*G*) Quantification of the cell fluorescence in (F). All the above experiments were repeated at least three times (means ± SDs; n = 3; Student’s *t-*test, *, *p* < 0.05; **, *p* < 0.01, ****, *p* < 0.0001). Scale bar: 10 μm.

PC6 also could detect the activity of SARM1 endogenously expressed in HEK293T cells (Zhao et al., 2019). CZ-48 activated the endogenous SARM1 and produced increase of cytosolic PAD6 signal (Fig. 2*D*, upper right), but none in the SARM1-knockout cells (Fig. 2*D*, right lower), confirming specificity.

HEK293-line carrying inducible SARM1 (Zhao et al., 2019) was used to further substantiate that the PAD6 fluorescence was derived from SARM1 activity. Without induction, only basal SARM1 (Figure 2—figure supplement 1*A*) with minimal activity was detected (Fig. 2*E*, green dots), which was activated by CZ-48, resulting in increase in PAD6-fluorescence (orange triangles). Induction of SARM1 (Figure 2—figure supplement 1*A*) produced minimal signal also (Fig. 2*E*, purple squares), confirming SARM1 is auto-inhibitory. With CZ-48, both the basal and the induced SARM1 were activated, resulting in the largest signal (Fig. 2*E*, black triangles). In SARM1-knockout cells, no signal was detected (Fig. 2*D*, SARM1-KO; Fig. 2*E*, blue and red dots).

Endogenous NMN can be increased by ablating NMN-adenylytransferase (NMNAT1)(Zhao et al., 2019), resulting in increased PAD6 fluorescence (Fig. 2*F*) in a time-dependent manner (Fig. 2*G*).

Consistent with the *in vitro* results, PC6 is highly selective for SARM1 over CD38 in live cells. Cells expressing either wildtype or Type III mutant CD38(Liu et al., 2017, Zhao et al., 2012) did not show PAD6-signal after 48-hour incubation with PC6 (Figure 2—figure supplement 1*B*), even though the expressed enzymes readily increased cellular cADPR (Figure 2—figure supplement 1*C*).

The online version of this article includes the following figure supplement for figure 2:

**Figure supplement 1.**
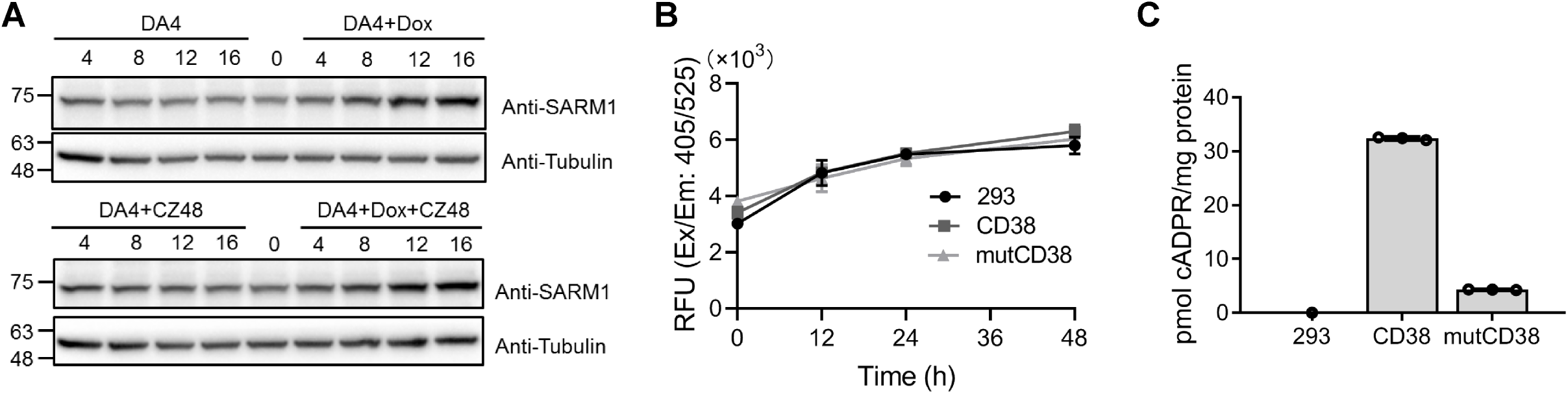
The expression level of SARM1 for fig.2E and the activities of CD38. (*A*) Western blots analysis of the expression of SARM1 in the inducible cell lines with or without treatment of 0.5 μg/ml Dox and 100 μM CZ-48 for the indicated time. (*B*) The HEK293T cells overexpressing wildtype and Type III mutant CD38 (mutCD38)(Liu et al., 2017, Zhao et al., 2012) were incubated with 50 μM PC6 for the indicated time and the fluorescence of PAD6 was analyzed in the flow cytometry; (*C*) The cellular cADPR levels in the cells (A) were measured by the cycling assay.(Graeff et al., 2002)

### Imaging SARM1 activation during AxD

Vincristine (VCR)-induced AxD in peripheral neuropathy is a common side effect of chemotherapy (Essuman et al., 2017) and is thought to be due to SARM1-activation (Gerdts et al., 2013). Mouse dorsal root ganglion (DRG) neurons were infected with lentivirus carrying TdTomato for visualizing the axons (Fig. 3*A*, 3*C*, orange), and with either a non-targeting (Fig. 3*A*) or SARM1-specific shRNA (Fig. 3*C*). In the non-targeting group, VCR elevated PAD6-fluorescence (Fig. 3*A*, green), indicating activation of SARM1, by as early as 4-8 hours and reaching a maximum by 16 hours (Fig. 3*A* and 3*D*, blue). AxD started at about 20 hours (Fig. 3*F*, blue; Figure 3—figure supplement 1A), temporally consistent with a causal role for SARM1. Another measure of SARM1 activation is the elevation of cellular cADPR (Zhao et al., 2019), which occurred (Fig. 3*E*, blue) by 12 hours, peaking at 24 hours. Neurons not treated with VCR showed neither SARM1-activation nor AxD (Fig. 3*A, D, E, F* and Figure 3—figure supplement 1*A*, CTRL).

**Fig. 3.**
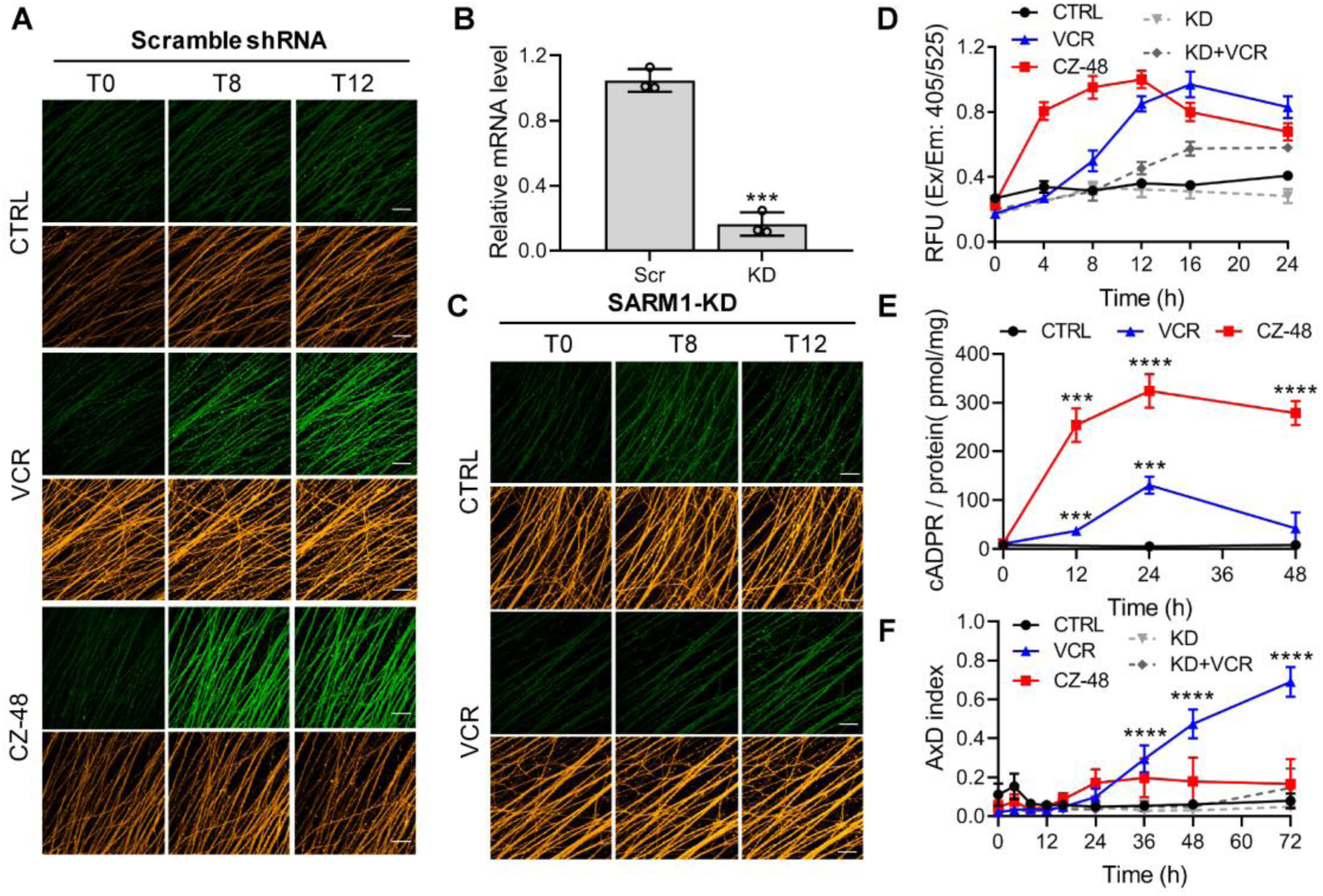
SARM1 activation in mouse DRG upon vincristine treatment. (*A,C*) Confocal imaging of SARM1 activation in DRG neuronal axons. The neurons were infected with virus expressing TdTomato to provide easy imaging of the axons. Cells were additionally transfected with either scramble (A) or SARM1-specific (C) shRNAs and treated with 50 μM PC6, 200 μM CZ-48 or 50 nM Vincristine and imaged in the indicated time points. Green: PAD4; Orange: TdTomato; scale bars: 50 μm. (*B*) Knockdown efficiency of SARM1. Scr, scramble shRNA; KD, SARM1-specific shRNA. (*D*) Quantification of the fluorescence intensity of PAD6 in DRGs. (*E*) Intracellular cADPR contents. (*F*) Indices of AxD. All the above experiments were repeated at least three times (means ± SDs; n = 3; Student’s *t-*test, ***, *p* < 0.001; ****, *p* < 0.0001).

Reducing endogenous SARM1 using shRNA (Fig. 3*B, D* and *F*, KD) reduced the PAD6 fluorescence without altering its peaking at 16 hours (Fig. 3*C*; 3*D*, KD+VCR) and reduced AxD (Fig. 3*F*, KD+VCR; Figure 3—figure supplement 1*B*), further substantiating a causal role for SARM1. CZ-48 induced SARM1-activation more rapidly (Fig. 3*A* and 3*D*, red) and elevated cADPR higher (Fig. 3*E*, red), confirming its direct action. Intriguingly, CZ-48 did not induce massive AxD as VCR (Fig. 3*F*, CZ-48; S5*A*). These results indicate SARM1-activation is a necessary and causal factor, but not a sufficient one for AxD. Other critical factors might be downstream events leading to microtubular dysfunction.

The online version of this article includes the following figure supplement for figure 3:

**Figure supplement 1.**
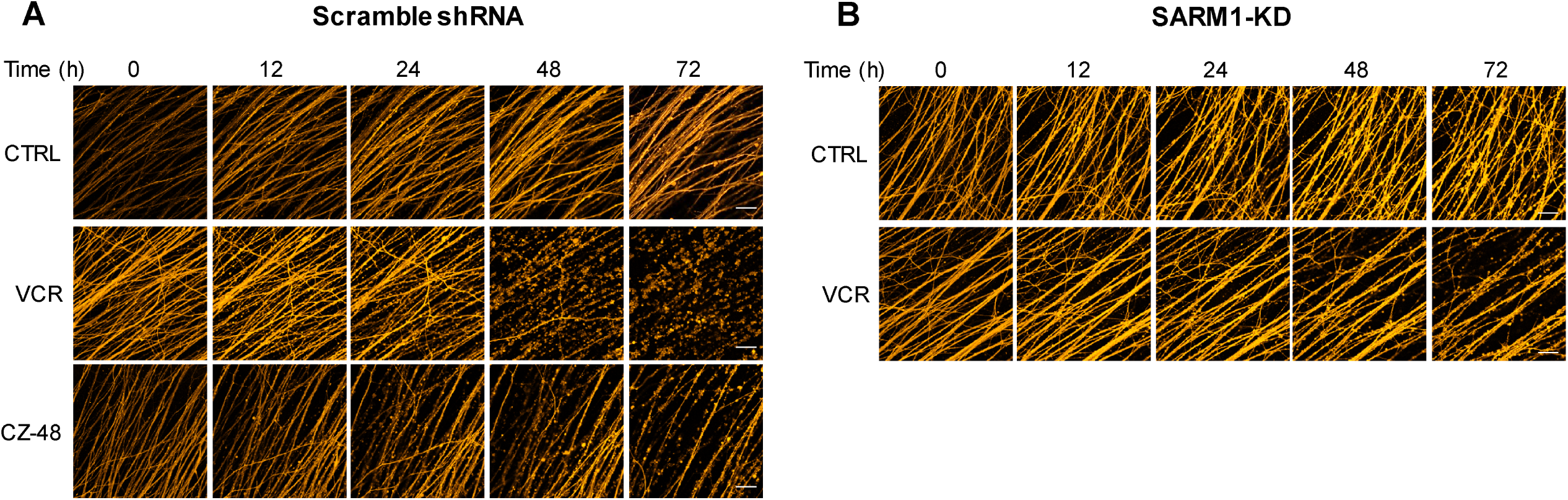
The integrity of axons visualized by the TdTomato fluorescence. The DRG neurons, on Div6, were infected with the lentivirus expressing TdTomato and scramble shRNA (*A*) or SARM1-specific shRNA (*B*) and 3 days later, incubated with 50 nM VCR or 200 μM CZ-48 with 50 μM PC6. The photos of axons with TdTomato fluorescence were captured under the confocal microscope (Nikon, A1). Scale bars: 50 μm.

### Dehydronitrosonisodipine (dHNN) is an inhibitor of SARM1 activation

Another prompt application of PC6 is library screening for inhibitors of SARM1. The feasibility was verified by measuring the IC_50_ of a reported inhibitor of SARM1, nicotinamide (Nam)(Essuman et al., 2017), to be around 140 μM (Figure 4—figure supplement 1*A*). NMN-activated SARM1 was incubated with drugs of the library (Fig. 4*A*) and its activity measured with PC6 in the presence of NAD (cf. Fig. 1). Out of 2015 drugs, 34 had more than 80% inhibition (Fig. 4*B*), which were further tested for inhibition of the SARM1-NADase activity using HPLC. Fig. 4*C* shows the plot the IC_50_-values of these drugs measured with both the PC6 and the NADase/HPLC assays. Twenty-four drugs are in the middle sector, indicating they inhibited both reactions similarly. Two inhibited the PC6 activity 5-fold less than the NADase (Fig. 4*C*, left sector), and eight in the right sector (7 have IC_50_s higher than 40 μM) inhibited NADase less than the base-exchange. These remarkable differences underscore the importance of using more than one assay for drug screening (see Discussion).

**Fig. 4.**
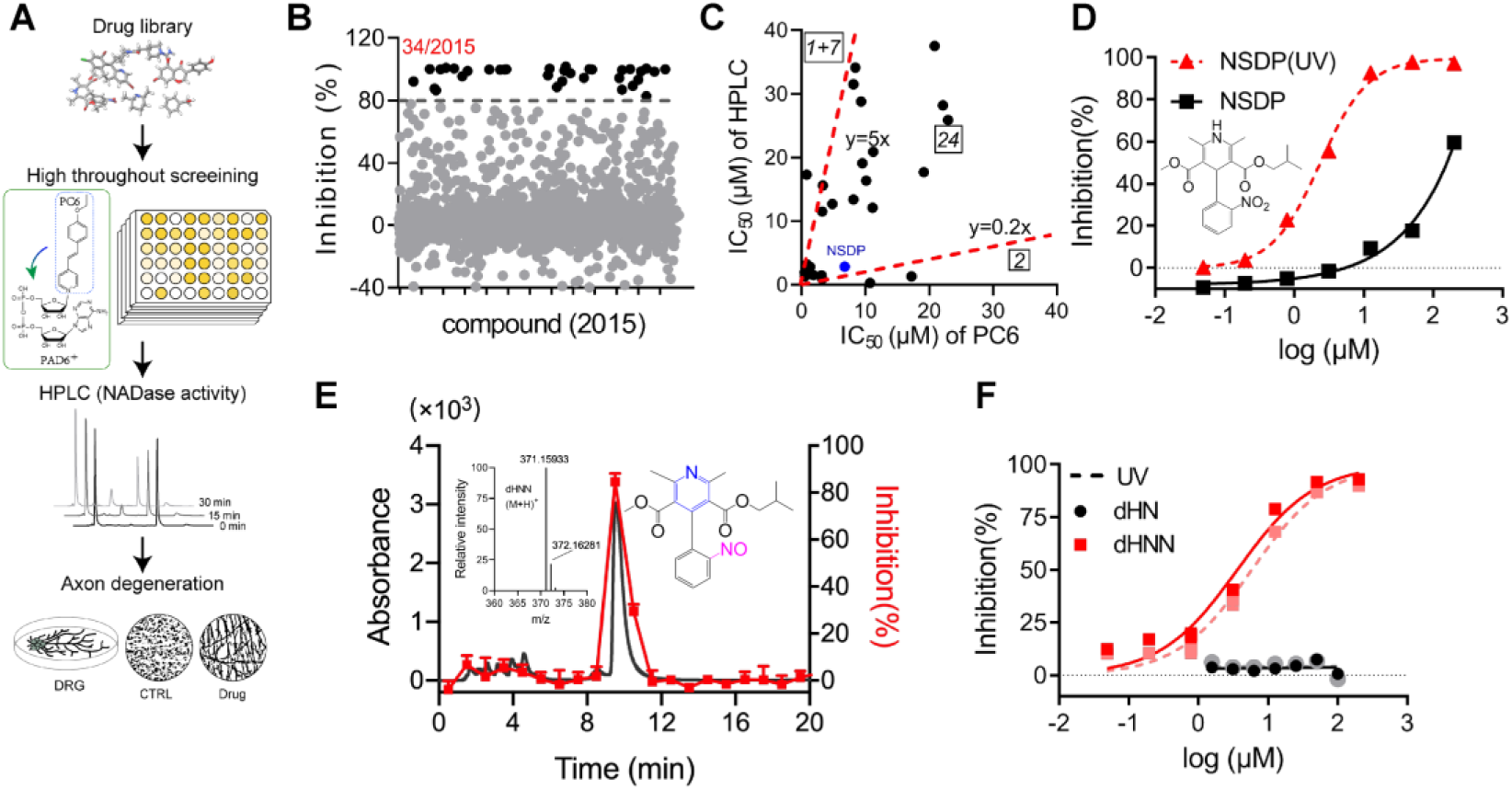
Identification of dHNN as an inhibitor of SARM1. (*A*) Flowchart of the PC6-based high-throughput screening. (*B*) Inhibitory effects of the 2,015 compounds (50 μM) from an approval drug library. The activity of drug-treated SARM1-dN was determined with PC6 assay. (*C*) Plot of IC_50_s of the 27 most potent inhibitory compounds from high-throughput screening, determined by PC6 (x axis) versus by HPLC (y axis) assays. See Method section. (*D*) Inhibition curves of NSDP before (Black) and after (NSDP(UV), red) UV at 254 nm for 30 min. (*E*) HPLC elution profile of dHNN. NSDP after 30 min UV treatment was analyzed using a C-18 column with a gradient of 0.1% TFA and ACN in 0.1% TFA. Fractions were assayed for inhibition of SARM1-dN by PC6 assay. The derivative in the elution peak was identified by MS. Black line: Absorbance at 275 nm; Red dots: inhibition activity. Insets: MS of the peak fraction showing its mass was the same as dHNN and the chemical structure of dHNN. (*F*) Concentration-inhibition curves of dHN (black solid line), UV-treated dHN (black dotted line), dHNN (red solid line) and UV-treated dHNN (red dotted line), measured by PC6 assay.

In the middle sector is Nisoldipine (NSDP), a calcium channel blocker having beneficial effects on neurodegenerative diseases. Peculiarly, its inhibition of SARM1 varied widely among batches. Investigations indicated the active compound was not NSDP but its derivative. Fig. 4*D* shows fresh NSDP had an IC_50_-value of about 150 μM (squares), but its potency increased 75-fold after exposure to UV (Fig. 4*D*, triangles, IC_50_=2 μM). Also, fresh NSDP had an HPLC-elution peak at 12.2 min (Figure 4—figure supplement 1*B*), but was completely converted by UV to a compound having a peak at 9.8 min that strongly inhibited of SARM1 (Fig. 4*E*, red). HRMS showed that the active compound had a mass of 370.15205 Da (Fig. 4*E*, inset) identical to a known derivative of NSDP, dehydronitrosonisoldipine (dHNN)(Marinkovic et al., 2003). The HPLC-elution profile of the active compound was also the same as dHNN (Figure 4—figure supplement 1*B*, purple line and green dash). Indeed, authentic dHNN was active and could not be further activated by UV (Fig. 4*F*, red line and dash). Another derivative of NSDP, dehydronisoldipine (dHN, elution peak at 8.7 min, Figure 4—figure supplement 1*B*), showed no inhibition before or after UV (Fig. 4*F*, black line and dash), indicating that the nitroso group is essential for the effect.

The online version of this article includes the following figure supplement for figure 4:

**Figure supplement 1.**
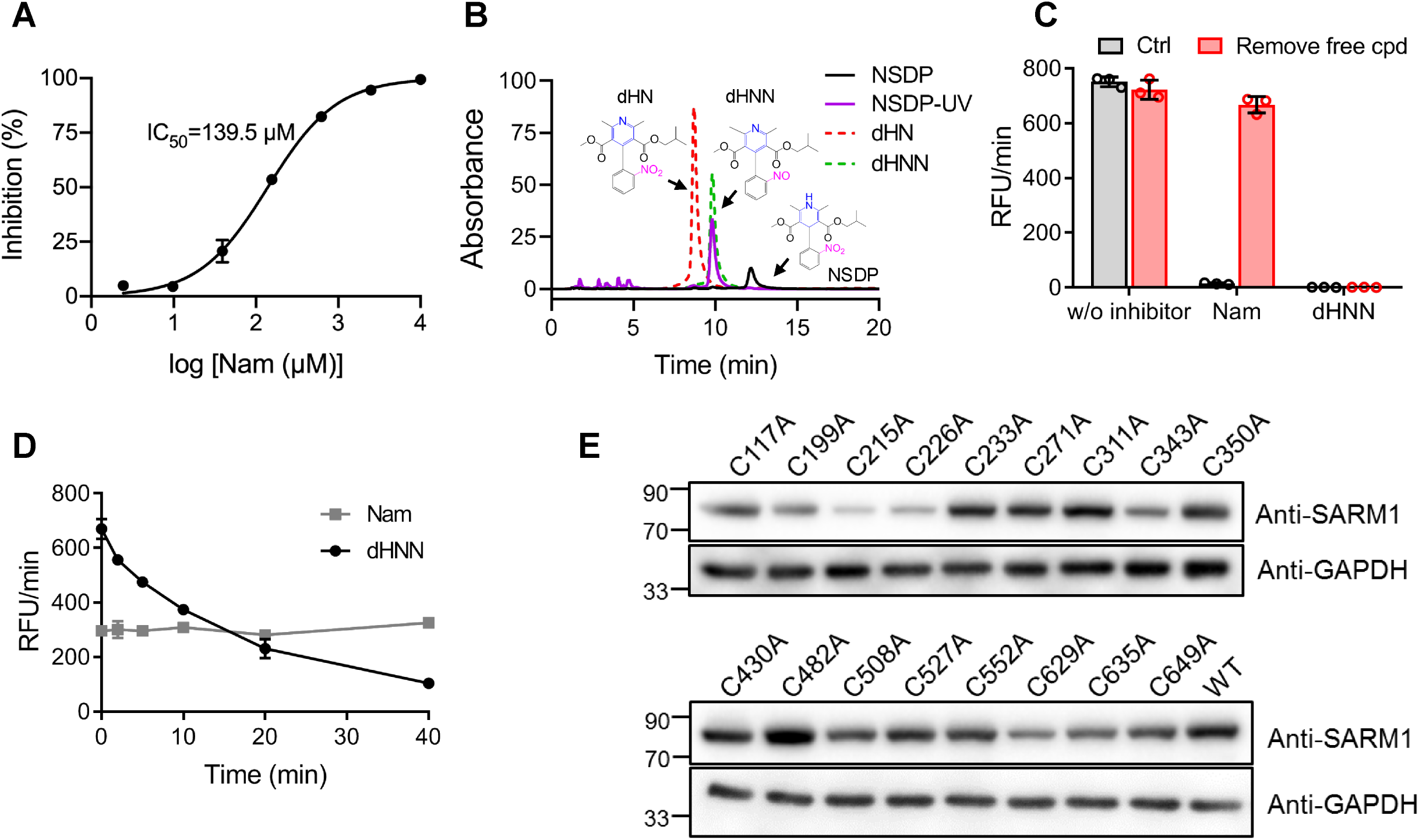
The inhibitory mechanism of dHNN against SARM1. (*A*) The inhibition curves of nicotinamide to SARM1-dN measured with PC6-based reaction. (*B*) HPLC analysis of NSDP after treating by UV at 254 nm for 30 min. black: standard of NSDP; purple: NSDP after UV; red: standard of dHN; green: standard of dHNN; (*C*) The irreversibility of the inhibition of dHNN. SARM1-dN was pre-incubated with 20 μM dHNN or 5 mM Nam at RT for 20 min and washed using Centricon filter. Activity was analyzed afterward with PC6 assay. (*D*) Time dependent inhibition of dHNN to SARM1-dN. 10 μM dHNN or 300 μM of Nam was pre-incubated with SARM1-dN for the indicated time. Activity was analyzed afterward with PC6 assay (*E*) Western blots analysis of the expression of SARM1 with single cysteine mutation to alanine. C215A and C226A show much lower protein expression, indicating the mutations seems affect the stability of the whole protein, although the IC_50_s of decreased to 30 μM and 11 μM, respectively, comparing to that of wildtype SARM1 (6 μM).

### dHNN inhibits SARM1 and AxD by covalently modifying cysteines

The dHNN-inhibition was irreversible by washing (Figure 4—figure supplement 1*C*, red bars), while that by Nam was reversible. Also, dHNN-inhibition was time-dependent, but not Nam (Figure 4—figure supplement 1*D*), strongly suggesting dHNN covalently reacted with SARM1.

To determine the target of dHNN, we truncated the inhibitory ARM-domain, producing a constitutively active SAM-TIR, which showed a right-shifted inhibition-curve comparing to the full-length form (Fig. 5*A*), with around 50-fold increase in the IC_50_. dHNN decreased the cellular cADPR in cells expressing SARM1, but not in those expressing SAM-TIR (Fig. 5*B*). These results suggest that dHNN is cell-permeant and acts mainly by blocking SARM1 activation and not its enzymatic activities.

**Fig. 5.**
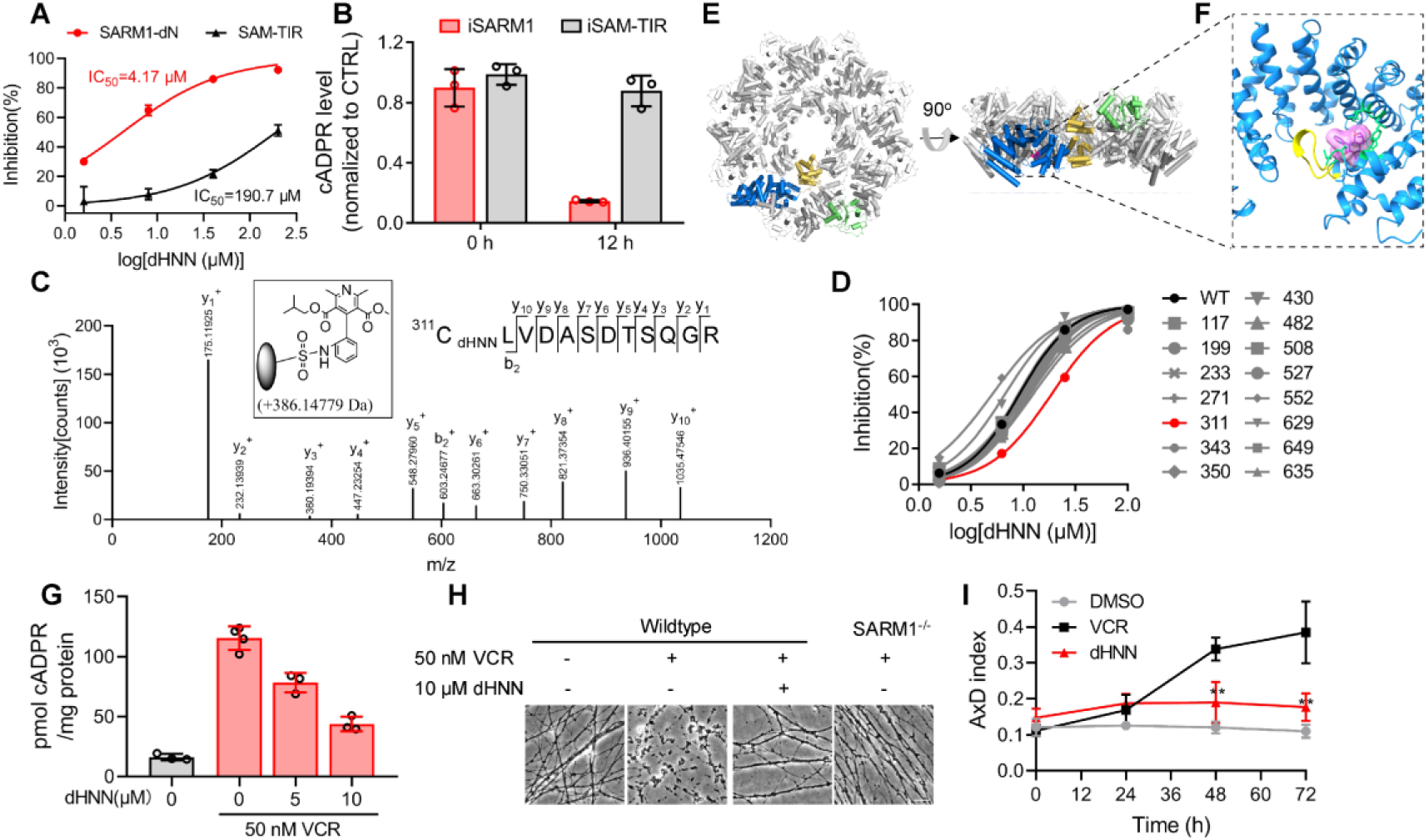
dHNN reduces AxD by inhibiting SARM1 through covalent modification of the cysteine. (*A*) Inhibition of SARM1-dN and SAM-TIR by dHNN *in vitro*. See Method. (*B*) Inhibition of SARM1-dN and SAM-TIR by dHNN *in cellulo*. See Method. (*C*) MS of SARM1-dN modification by dHNN. Peptide spectrum match shows that Cys311 was modified by dHNN, increasing its mass to 386.1483 Da. (*D*) Each cysteine in SARM1-dN was mutated to alanine. The dHNN-IC_50_s were measured by PC6 assay. (*E*) Top (left) and side (right) view of the SARM1 octamer. α-helices are shown as cylinders. dHNN modifications are shown as sticks. One protomer is colored in blue for ARM, gold for SAM and green for TIR. The other protomers are colored in grey. (*F*) Zoom-in view of the dHNN-modified pocket. dHNN: purple stick and molecular surface; interacting residues: green; loop: yellow. (*G*) DRG neurons were treated with dHNN for 16 h in the presence of VCR. The cellular cADPR contents were measured. (*H*) Micrographs of AxD of DRG neurons after treatment of VCR in presence of dHNN for 72 h. (*I*) Quantification of AxD indices after 0, 24, 48, 72 h treatment with VCR as in (H).

The nitro group of dHNN may covalently modify cysteines (Callan et al., 2009) in SARM1. Indeed, LC-MS/MS identified dHNN-modifications (+386.1483 Da, Fig. 5*C*, inlet) were mainly in Cys311 of the ARM domain (Fig. 5*E*, Figure 5—figure supplement 1*A*-*L*). Many peptides of other proteins with cysteines were also identified but none showed modification by dHNN (Figure 5—figure supplement 1*M*), indicating specificity of dHNN. Single mutation of all the cysteines to alanine showed that Cys311A significantly decreased the response to dHNN (Fig. 5*D*).

With Cryo-EM, we found that dHNN stabilized a similar inhibitory conformation of SARM1 as that induced by NAD (PDB: 7cm6)(Jiang et al., 2020). In 2D-classification of the untreated SARM1, most particles presented only the SAM octamer ring (Figure 5—figure supplement 3*A*). For the dHNN-treated SARM1, larger octamer ring corresponding to both the SAM and ARM/TIR domains could be clearly observed (Figure 5—figure supplement 2, Supplementary file 1, Figure 5—figure supplement 3*B*). Its 3D-structure was constructed at 2.4 Å resolution (Figure 5—figure supplement 3*C*) with residues from 56 to 549 (ARM and SAM domains) and 561 to 702 (TIR domain) all fitted into the cryo-EM map (Fig. 5*E* and Figure 5—figure supplement 3*D*). Structural superimposition with the NAD-bound SARM1 (PDB: 7cm6)(Jiang et al., 2020) showed RMSD values of 0.91 (Figure 5—figure supplement 3*E*), suggesting that dHNN constrains SARM1 in an inactive conformation similar to that induced by NAD. Extra electron density was only observed near residue Cys311 (Figure 5—figure supplement 4*A*), the dHNN-target (Fig. 5*C*, 5*D*), but not other cysteines, consistent with it being derived from dHNN (5*F*, purple). dHNN interacts with Leu265, Leu268, Phe308, Arg322 and Tyr348 (Fig. 5*F* and Figure 5—figure supplement 4*A*, green) in the ARM domain, pushing the insertion loop (Fig. 5*F*, yellow) towards ARM1 and stabilizes the domain. This is similar to that observed with NAD, which binds at the other side of the insertion loop (Figure 5—figure supplement 4*B*) and stabilizes the ARM domain possibly via ligating ARM1 and the insertion loop (Figure 5—figure supplement 4*B*).

By preventing SARM1 from activation, dHNN also inhibited the VCR-activated cADPR production (Fig. 5*G*) in neurons and blocked not only the VCR-induced AxD (Fig. 5*H*, 3^rd^ picture; Fig. 5*I*, red line) but also AxD after axotomy (Figure 5—figure supplement 5*A*, 3^rd^ column; *B*, red line) as effective as knocking out SARM1 (Figure 5—figure supplement 5*B*, grey lines).

The online version of this article includes the following figure and file supplements for figure 5:

**Figure supplement 1.**
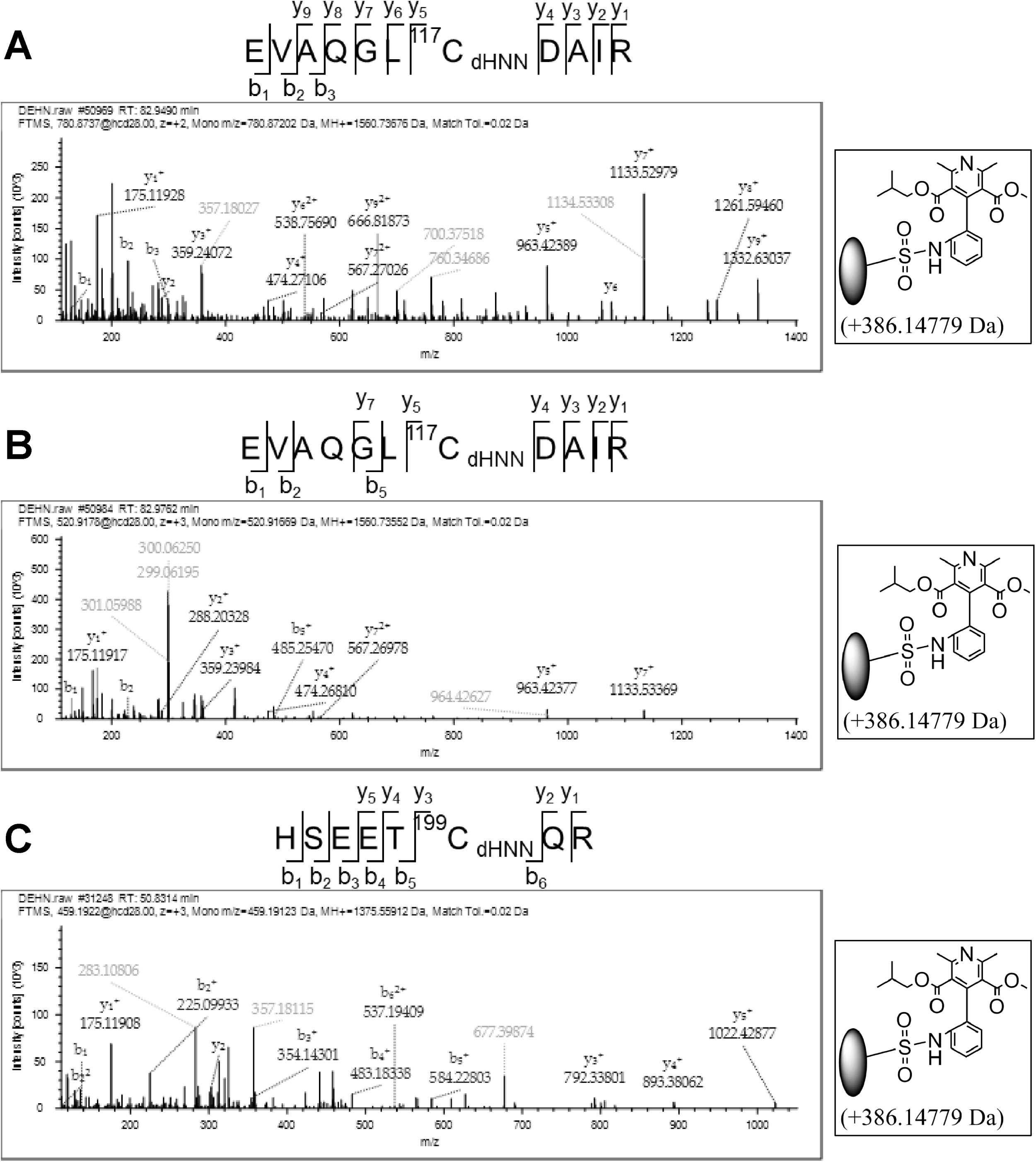
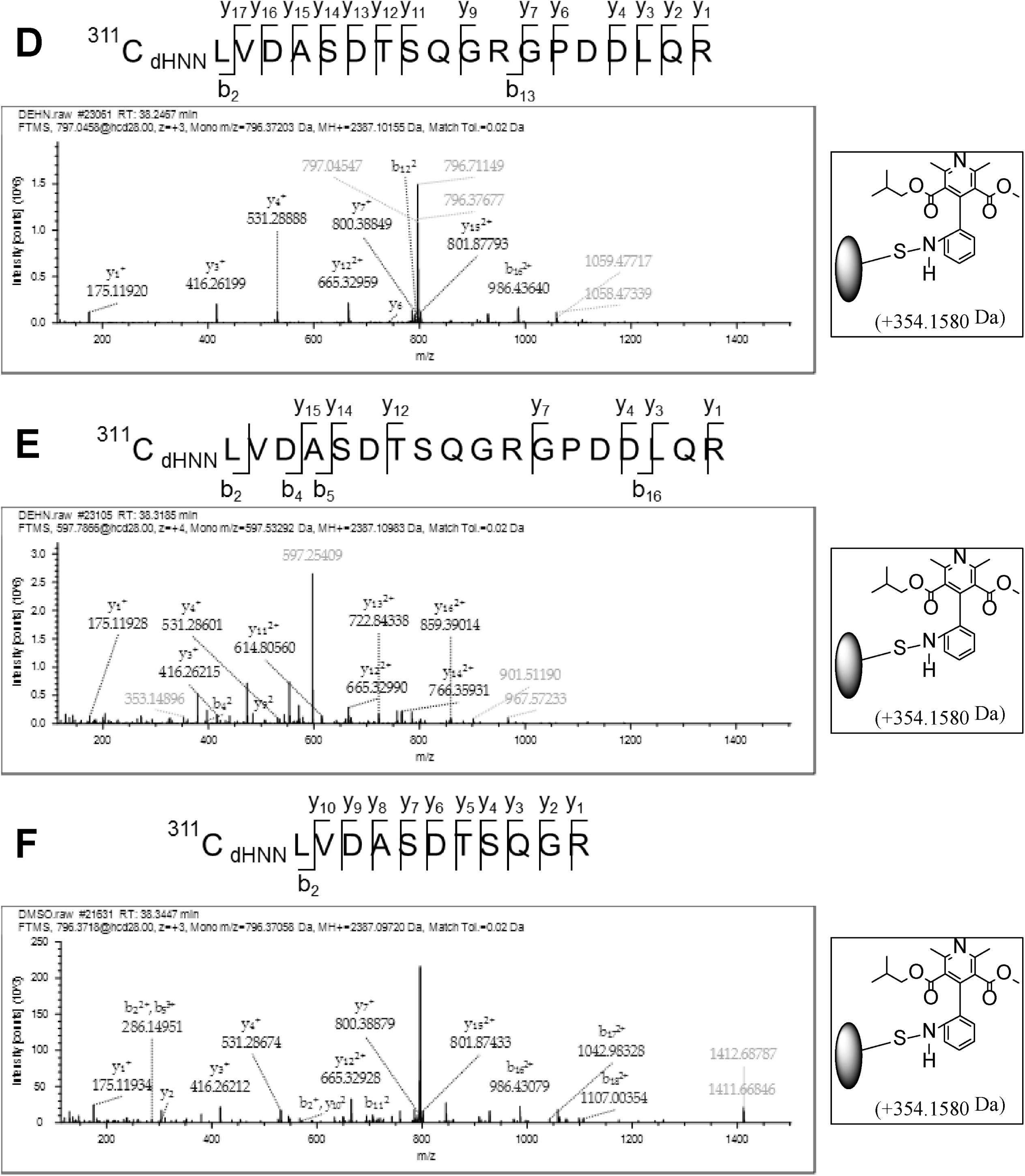
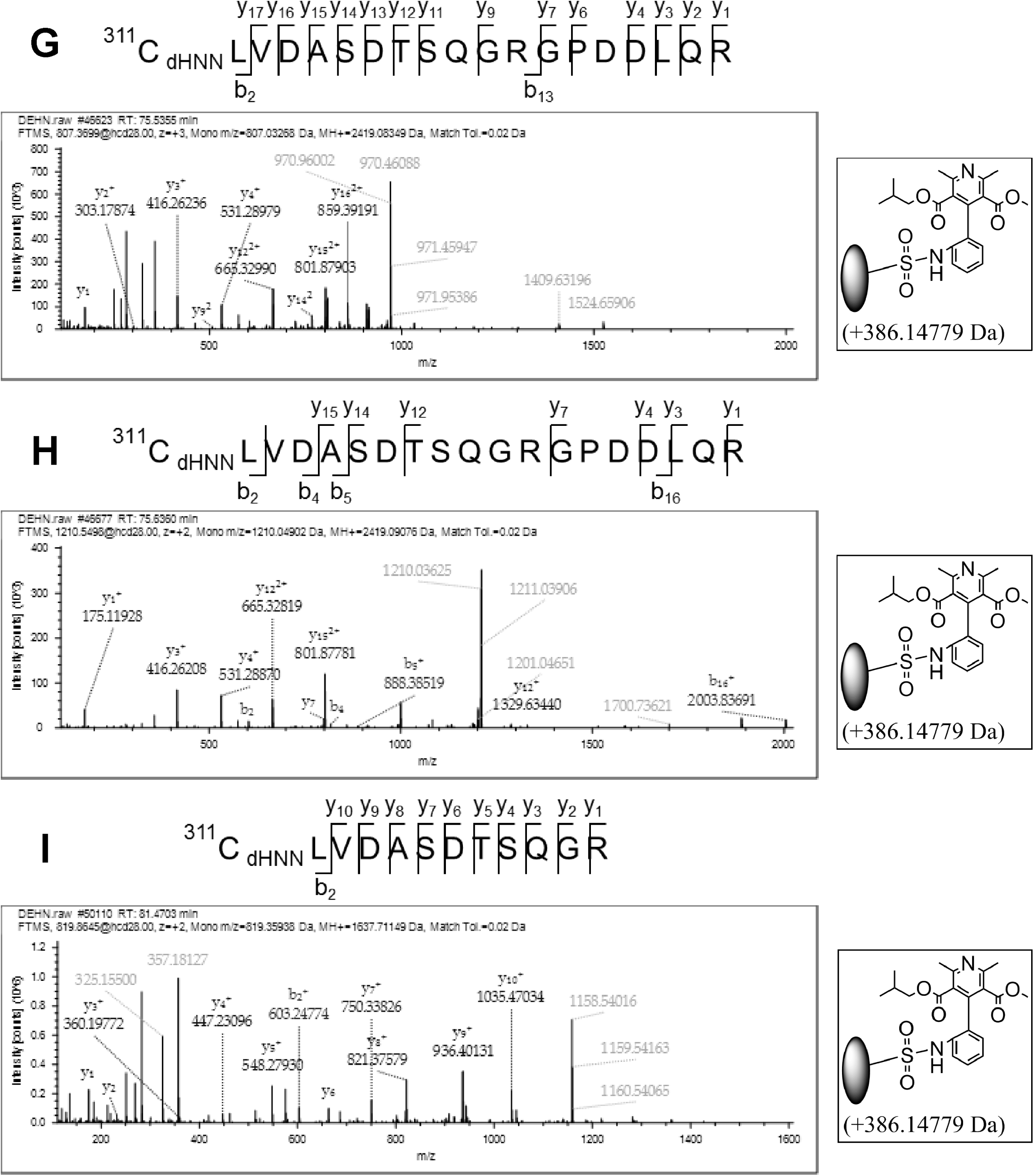
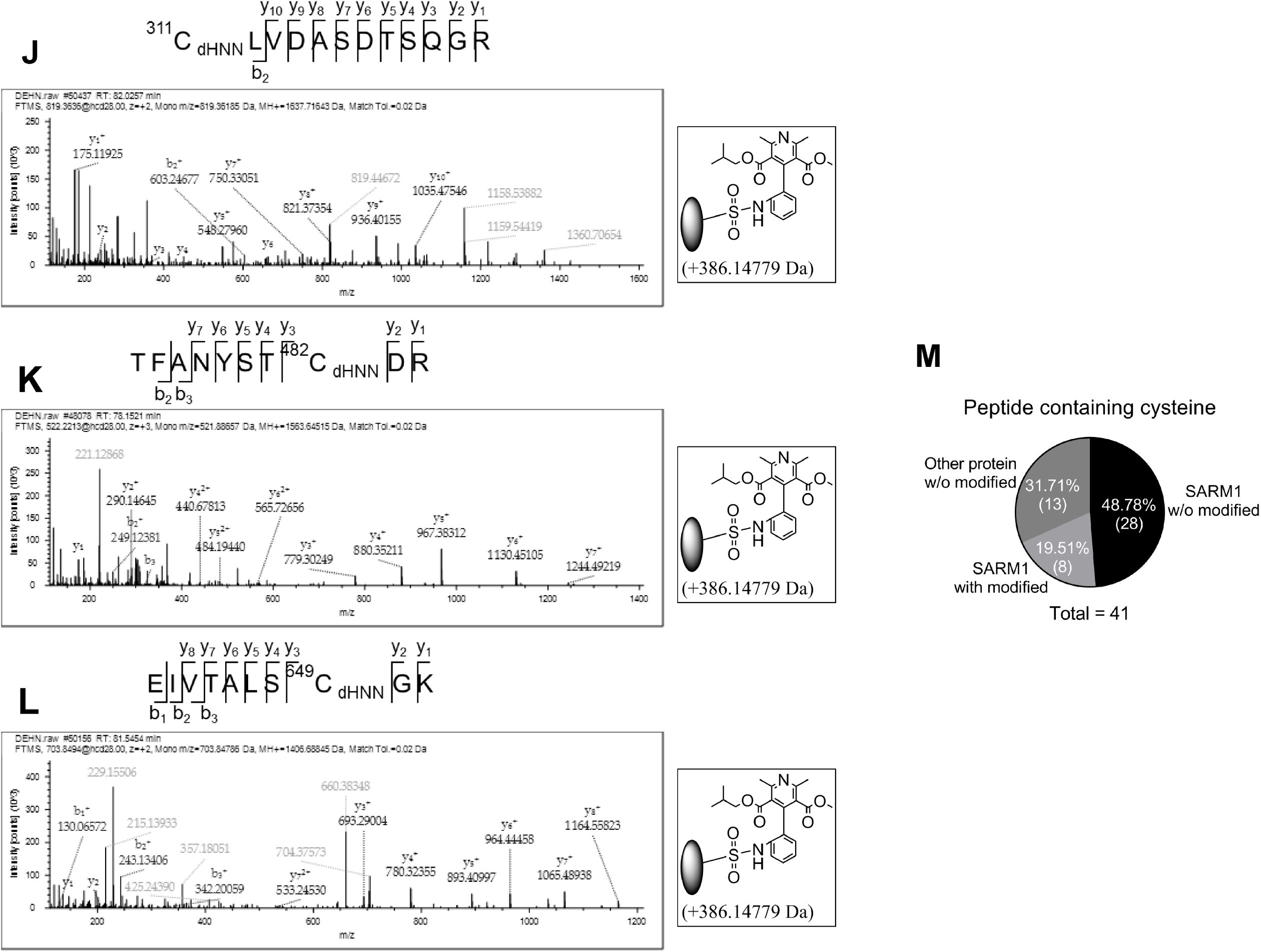
The dHNN modifications on the peptides of SARM1 analyzed by LC-MSMS. (*A-L*) The peptides of SARM1 containing the dHNN modifications characterized by LC-MSMS. (M) The statistical analysis of the peptides containing cysteines with or without the dHNN modifications characterized by LC-MS/MS.

**Figure supplement 2.**
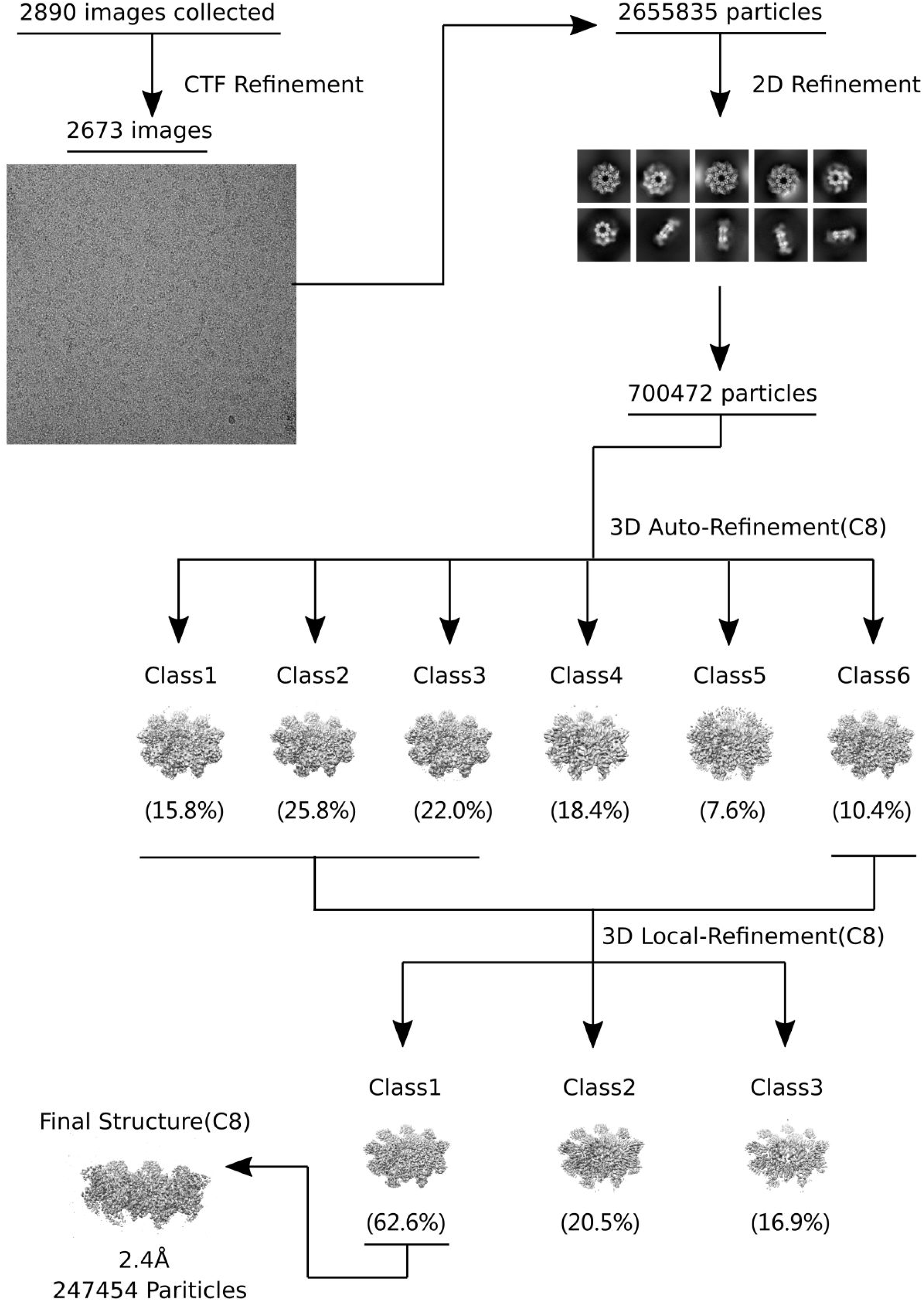
Data processing procedure for the SARM1-dHNN structure.

**Figure supplement 3.**
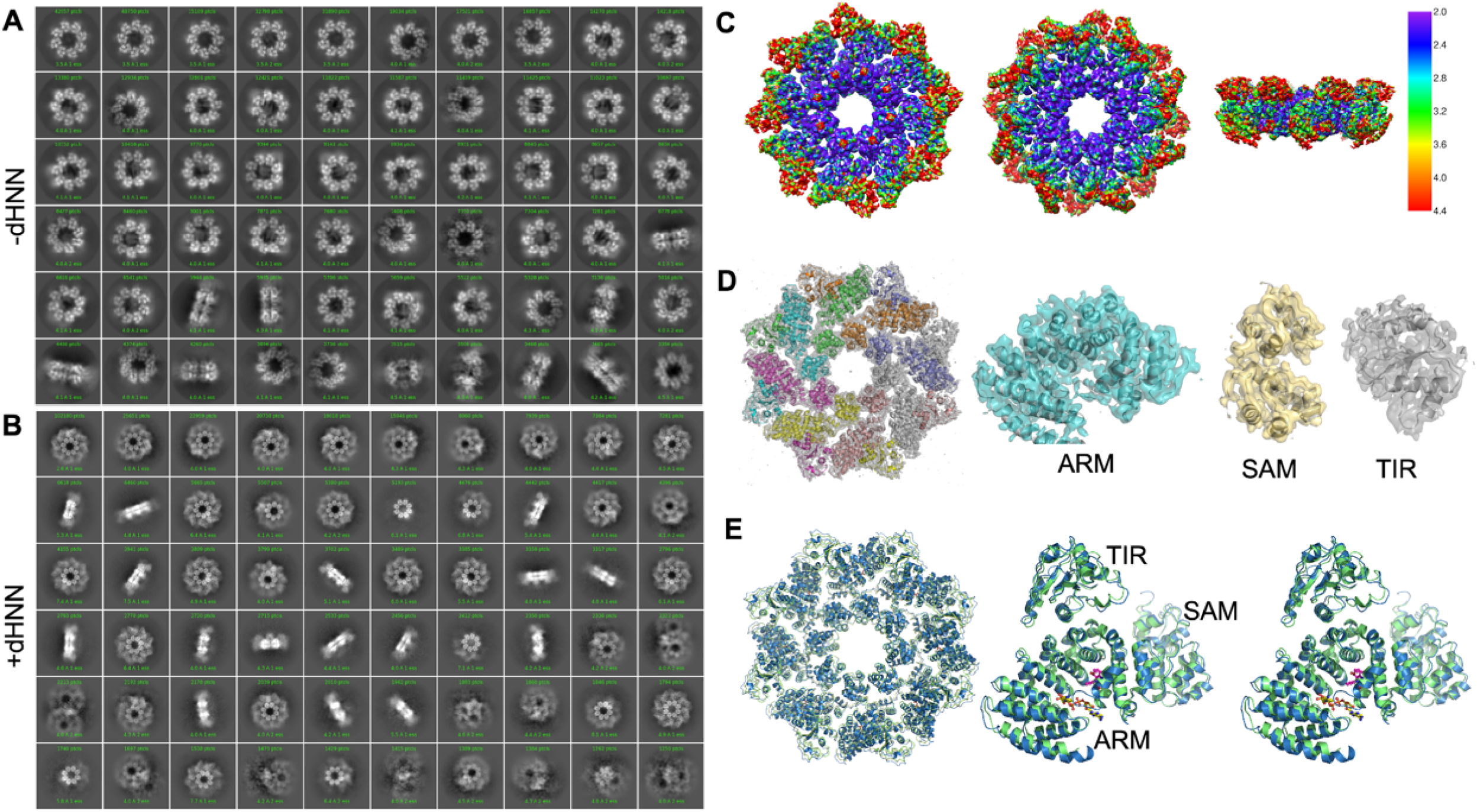
The structure of SARM1 was stabled in inactive form after dHNN treatment. (*A-B*) Representative 2D class averages of SARM1-dN in the absence (*A*) or presence (*B*) of 50 μM dHNN in 100 mM Tris, 150 mM NaCl and 1 mM EDTA at pH 8.0. (C) Local resolution estimation calculated by RELION for the 3D reconstruction of SARM1-dHNN. Left, top view; middle, bottom view; right, side view. (*D*) Density fit to the 3D reconstruction. (*E*) Superimposition of SARM1-dHNN onto SARM1-NAD (PBD codes 7cm6 and 7cm7) shows no obvious structural deviation. Left: octamer, blue for SARM1-dHNN, green for 7cm6 and grey for 7cm7; right, stereo view for the superimposition of one protomer in SARM1-dHNN and SARM-NAD (7cm6).

**Figure supplement 4.**
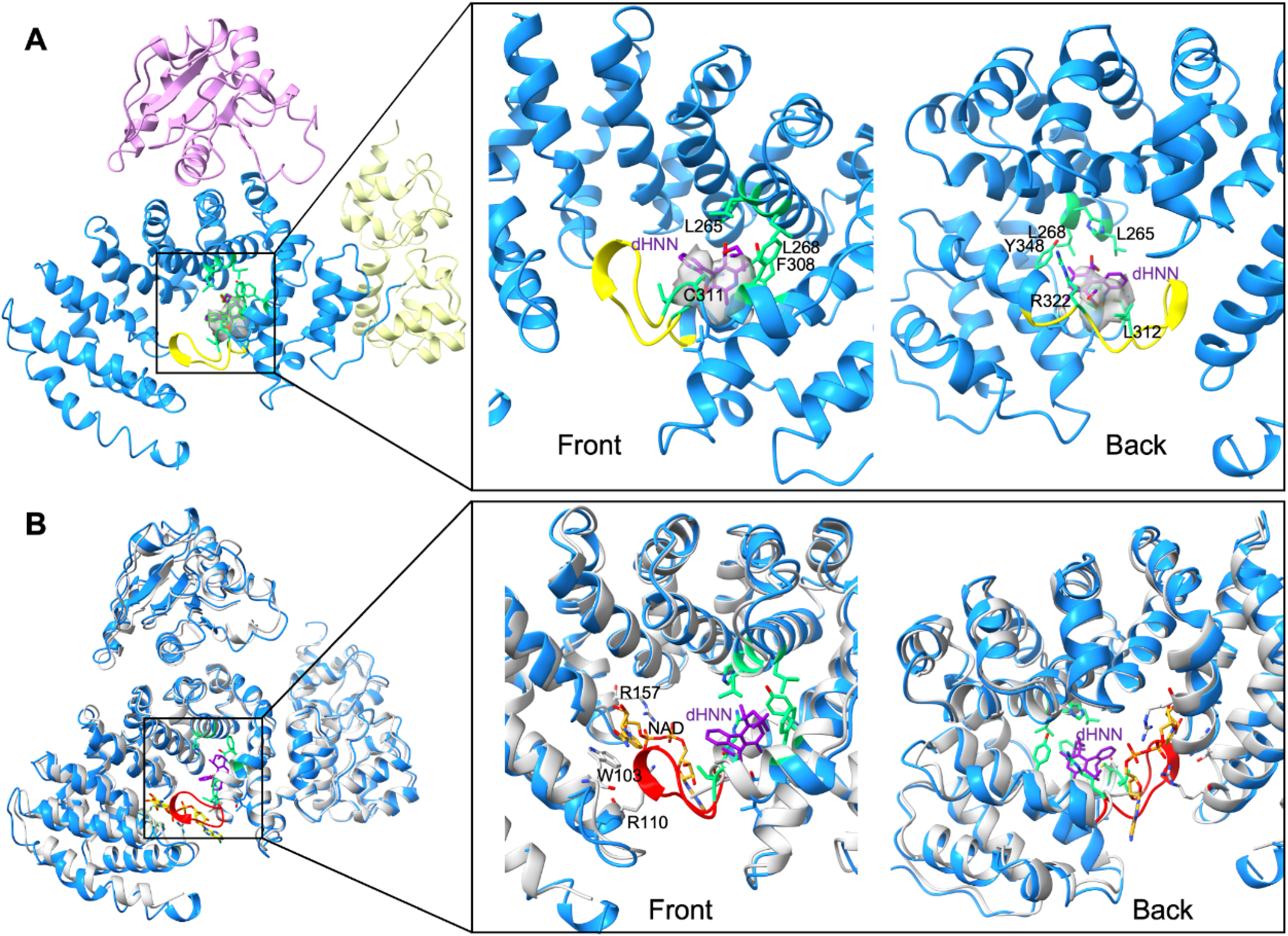
dHNN bound to residue C311 in the ARM domain of SARM1 (*A*) dHNN covalently links to residue C311 in ARM domain. ARM, SAM and TIR domains were shown as cartoon and colored in blue, yellow, and pink, respectively. The insertion loop between residues L312 and G323 was colored in yellow. The electron density corresponding to dHNN was shown as grey surface. dHNN and residues interacting with dHNN were shown as stick models and were colored in violet and green, respectively. (*B*) Superposition of SARM1-dHNN onto SARM1-NAD (PDB code 7mc6). SARM1-dHNN and SARM1-NAD were shown as blue and grey cartoon, respectively. The insertion loop was in SARM1-dHNN was shown in red. NAD was shown as stick models and colored with gold carbons. Residues interacting with NAD, W103, R110 and R157, were shown as stick models with grey carbons. dHNN and residues interacting with dHNN were shown as in (A).

**Figure supplement 5.**
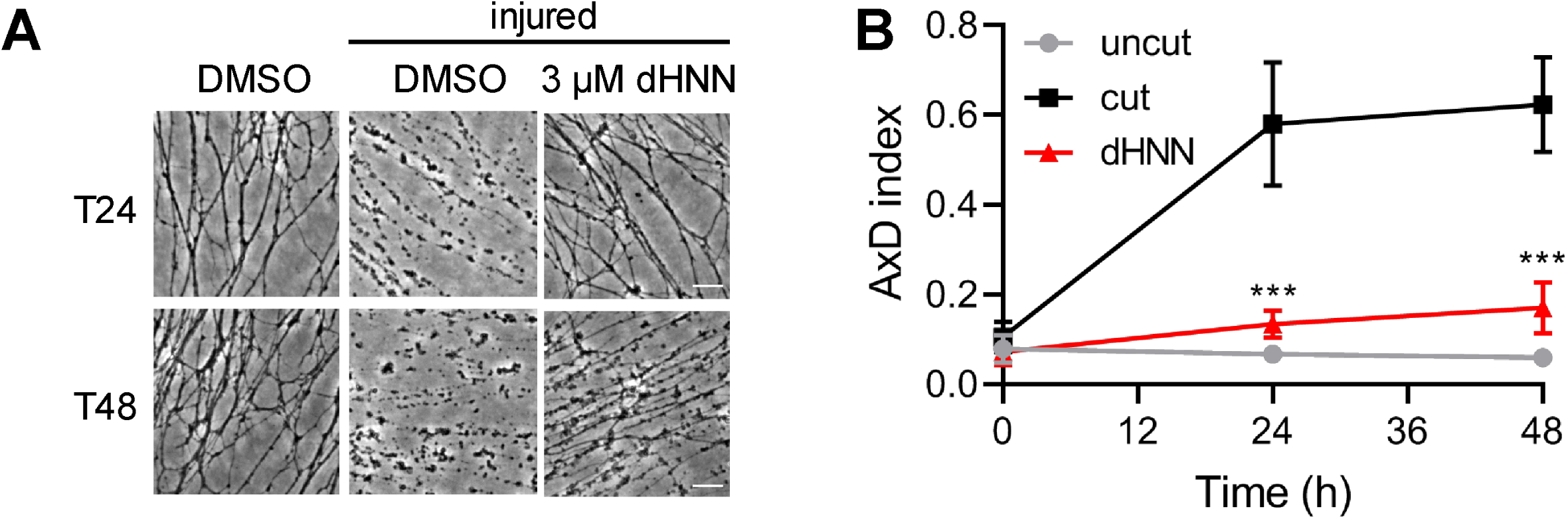
dHNN attenuates the axotomy-induced AxD. DRG neurons were pre-treated with 3 μM dHNN for 0.5 h and axotomy performed to induce AxD. Images were capture at 0, 24, 48 h (*A*) and the AxD index was analyzed by ImageJ (*B*).

**Supplementary file 1**. Refinement statistics for the SAMR1-dHNN structure

## Discussion

Visualizing the activity of a signaling enzyme in live cells provides clearer understanding of the spatial and temporal aspects of its mechanism and function, a goal sought by many. The PC-probes developed here are particularly advantageous. They are cell permeant, but the SARM1-catalyzed exchange products are impermeant and accumulates in the cytosol, enhancing its detection. The remarkably large red shift of the product fluorescence provides easy visualization away from the interference of auto-fluorescence. Our approach requires neither expression of construct nor cell manipulation. That it is applicable to any live cells was documented using CZ-48, a cell permeant activator of SARM1. With the probes, we provided the first direct evidence in live DRG neurons that SARM1 activation precedes AxD by several hours and that it is a necessary but insufficient factor for AxD.

Screening library to identify compounds of interest is a straightforward strategy widely used. The case for SARM1 is more complicated, as it is not only a multi-domain protein but also an auto-regulated enzyme catalyzing multiple reactions. Compounds may target the regulatory ARM domain as shown here for dHNN, or the catalytic TIR domain as Nam thought to be. For SARM1, the substrates are different for the base-exchange and the NADase reactions and may thus be differentially affected by the inhibitor-induced conformational changes of the catalytic site. Although the exact reason remains to be determined, the compounds shown here that can selectively block one reaction much more than the other are of interest. Many believe that the NADase activity of SARM1, leading to cellular NAD depletion, is its dominant property for regulating AxD. But the two calcium messengers, cADPR and NAADP produced by its cyclase and base-exchange reactions may well have functional roles as well. Compounds with differential inhibition can thus be an important tool to resolve the issue.

Much effort is being invested in targeting SARM1-mediated NAD depletion for therapeutic protection from AxD. Chemical blockers may well be an ideal tool for turning off the NAD depletion. dHNN uncovered in this study is the first compound ever described that can block the activation of SARM1, revealing a druggable allosteric site and can thus usher in a new approach for therapeutic drug development.

Another point of interest is that dHNN is a derivative of NSDP. Metabolic conversion of NSDP to dHNN, leading to inhibition of SARM1, may well account for the neural protective effects of NSDP (Siddiqi et al., 2019). That NSDP is already a clinical drug makes the path of dHNN to therapeutic use smoother.

## Materials and Methods

### Animals

This study was carried out in strict accordance with animal use protocol approved by Peking University Shenzhen Graduate School Animal Care and Use Committee (#AP0015001). All animals (C57BL6/J), purchased from Guangdong Medical Laboratory Animal Center (China), were handled in accordance with the guidelines of the Committee on the Ethic of Animal Experiments. All surgery was performed after euthanasia and efforts were made to minimize suffering.

### Reagents

NAD, NMN, Digitonin, Poly-L-lysine, 5-fluoro-2’-deoxyuridine and uridine, KH_2_PO_4_, (NH4)_2_SO_4_ and iodoacetic acid were purchased from Sigma-Aldrich. DMEM, Neurobasal™ Plus Medium, Trypsin-EDTA, penicillin/streptomycin solution, B27 plus, GlutaMax, Laminin, Lipofectamine 2000, ConA-Alex-647, formic acid, Acetontrile were purchased from Thermo Fisher. NGF was purchased from Sino Biological. FBS was obtained from PAN Bitotech. Approval drug library (L1000) and Nisoldipine power (Cas # 63675-72-9) were purchase from Targetmol. Dehydro Nisoldipine (Cas #103026-83-1) was obtain from TRC while dehydronitrosonisoldipine (Cas #87375-91-5) were purchase from Glpbio and TRC. Vincristine was purchase from Selleck. General chemicals for probe synthesis were purchased from Dieckmann, Alfa, Energy or Sangon Biotech (Shanghai).

### Synthesis and characterization of pyridine conjugates (PCs)

All air and water sensitive reactions were carried out with anhydrous solvents in flame-dried flasks under argon atmosphere, unless otherwise specified. All the reagents were obtained commercially and used without further purification, unless otherwise specified. Anhydrous DMF was vacuum distilled from barium oxide, acetonitrile and dichloromethane was distilled from calcium hydride. Yields refer to isolated yields, unless otherwise specified. Reactions were monitored by thin-layer chromatography (TLC) carried out on 0.25 mm silica gel plates (60F-254) that were analyzed by UV light as visualizing method and by staining with anisaldehyde (450 mL of 95% EtOH, 25 mL of conc. H_2_SO_4_, 15 mL of acetic acid, and 25 mL of anisaldehyde) or KMnO_4_ (200 mL H_2_O of 1.5 g KMnO_4_, 10 g K_2_CO_3_ and 1.25 mL of 10% aq. NaOH). Silica gel (200-300 mesh) was used for flash column chromatography. Nuclear magnetic resonance (NMR) spectra were recorded on either a 300 (^1^H: 300 MHz, ^13^C: 75 MHz), 400 (^1^H: 400 MHz, ^13^C: 100 MHz), or 500 (^1^H: 500 MHz, ^13^C: 125 MHz) NMR spectrometer. The following abbreviations were used to explain the multiplicities: s = singlet, d = doublet, t = triplet, q = quartet, dd = doublet of doublets, m = multiplet, br = broad. High resolution mass spectra (HRMS) were obtained from a MALDI-TOF mass spectrometer.

To synthesize PC1, to a stirred solution of 4-vinylpyridine (210 mg, 2.0 mmol), 4-iodoaniline (220 mg, 1.0 mmol), P(*o*-tol)_3_ (61 mg, 20 mol%) and triethylamine (0.40 mL, 2.9 mmol) in degassed CH_3_CN (15 mL) under argon was added Pd(OAc)_2_ (23 mg, 10 mol%) quickly. The resulting mixture was stirred at 100 °C for 5 h. The mixture was then diluted with water (20 mL) and aqueous phase was extracted with ethyl acetate (15 mL × 3). The combined organic extracts were dried over anhydrous Na_2_SO_4_, filtered and evaporated under reduced pressure. Silica gel flash column chromatography (ethyl acetate/hexanes = 3:1) of the residue gave a pale-yellow solid (66 mg, 34%) as the product. PC1: mp = 272-273 °C. ^1^H NMR (400 MHz, DMSO-*d*_6_) δ 8.47 (d, *J* = 5.3 Hz, 2H), 7.46 (d, *J* = 6.0 Hz, 2H), 7.42 – 7.31 (m, 3H), 6.88 (d, *J* = 16.4 Hz, 1H), 6.59 (d, *J* = 8.5 Hz, 2H), 5.51 (s, 2H). ^13^C NMR (100 MHz, DMSO-*d*_6_) δ 149.8, 145.4, 133.7, 128.5, 123.6, 120.0, 120.0, 113.8. HRMS (+ESI) m/z calcd. for C_13_H_12_N_2_ (M+H)^+^ 197.1073, found 197.1072.

To synthesize PC2, to a stirred solution of 4-iodoaniline (0.60 g, 2.7 mmol) in DMF (7.5 mL) was added ethyl bromide (0.25 mL, 3.35mmol) and Na_2_CO_3_ (0.50 g, 4.72 mmol) at rt. The resulting mixture was stirred at 70 °C for 6 h. The mixture was then diluted with water (20 mL) and the aqueous phase was extracted with ethyl acetate (15 mL × 3). The combined organic extracts were washed with water (15 mL × 3), dried over anhydrous Na_2_SO_4_, filtered and evaporated under reduced pressure. Silica gel flash column chromatography (ethyl acetate/hexanes = 1:20) gave a brown solid (79 mg, 12%) as the product (**1**). Spectral data of **1** is consistent with those reported in the literature.^(Ni et al., 2017)^ To a stirred solution of 4-vinlypridine (53 mg, 0.5 mmol), **1** (74 mg, 0.3 mmol), P(*o*-tol)_3_ (30 mg, 20 mol%) and triethylamine (0.40 mL, 2.9 mmol) in degassed CH_3_CN (5 mL) under argon was added Pd(OAc)_2_ (11 mg, 10 mol%) quickly. The resulting mixture was stirred at 100 °C for 12 h. The mixture was then diluted with water (20 mL) and aqueous phase was extracted with ethyl acetate (15 mL × 3). The combined organic extracts were dried over anhydrous Na_2_SO_4_, filtered and evaporated under reduced pressure. Silica gel flash column chromatography (ethyl acetate/hexanes = 2:1) of the residue gave a pale-orange solid (51 mg, 76%) as the product. PC2: mp = 199-200 °C. ^1^H NMR (400 MHz, CDCl_3_) δ 8.52 (d, *J* = 5.8 Hz, 2H), 7.39 (d, *J* = 8.5 Hz, 2H), 7.31 (dd, *J* = 4.7, 1.4 Hz, 2H), 7.23 (d, *J* = 16.3 Hz, 1H), 6.79 (d, *J* = 16.2 Hz, 1H), 6.60 (d, *J* = 8.6 Hz, 2H), 3.21 (q, *J* = 7.1 Hz, 2H), 1.29 (t, *J* = 7.2 Hz, 3H). ^13^C NMR (125 MHz, CDCl_3_) δ 149.9, 149.0, 145.5, 133.4, 128.5, 125.2, 121.2, 120.4, 112.6, 38.2, 14.8. HRMS (+ESI) m/z calcd. for C_15_H_16_N_2_ (M+H)^+^ 224.1386, found 224.1382.

To synthesize PC3, to a stirred solution of 4-iodoaniline (1.2 g, 5.5 mmol) in DMF (15 mL) was added ethyl bromide (2.0 mL, 27 mmol) and Na_2_CO_3_ (1.0 g, 9.4 mmol) at rt. The resulting mixture was stirred at 70 °C for 6 h. The mixture was the diluted with water (20 mL) and the aqueous phase was extracted with ethyl acetate (15 mL × 3). The combined organic extracts were washed with water (15 mL × 3), dried over anhydrous Na_2_SO_4_, filtered and evaporated under reduced pressure. Silica gel flash column chromatography of the residue (ethyl acetate/hexanes = 1: 30) gave a brown oil (814 mg, 54%) as the product (**2**). Spectral data of **2** is consistent with those reported.^(Kolvari et al., 2014)^ To a stirred solution of **2** (273 mg, 1.0 mmol), 4-vinylpyridine (210 mg, 2.0 mmol), P(*o*-tol)_3_ (61 mg, 20 mol%) and triethylamine (0.40 mL, 2.9 mmol) in degassed CH_3_CN (15 mL) under argon was added Pd(OAc)_2_ (23 mg, 10 mol%) quickly. The resulting mixture was stirred at 100 °C for 12 h. The mixture then was diluted with water (20 mL) and the aqueous phase extracted with ethyl acetate (15 mL × 3). The combined organic extracts were dried over anhydrous Na_2_SO_4_, filtered and evaporated under reduced pressure. Silica gel flash column chromatography (ethyl acetate/hexanes = 1:1) of the residue gave a pale-yellow solid (138 mg, 55%) as the product. **PC3**: mp = 184-185 °C. ^1^H NMR (400 MHz, CDCl_3_) δ 8.50 (d, *J* = 5.6 Hz, 2H), 7.47 – 7.38 (m, 2H), 7.30 (d, *J* = 6.1 Hz, 2H), 7.25 – 7.19 (m, 1H), 6.74 (t, *J* = 16.9 Hz, 1H), 6.66 (t, *J* = 5.9 Hz, 2H), 3.47 – 3.18 (m, 4H), 1.31 – 1.02 (m, 6H). ^13^C NMR (100 MHz, CDCl_3_) δ 149.9, 148.2, 145.7, 133.4, 128.6, 123.2, 120.4, 120.4, 111.5, 44.5, 12.7. HRMS (+ESI) m/z calcd. for C_17_H_20_N_2_ (M+H)^+^ 253.1699, found 253.1699.

To synthesize PC4, to a stirred solution of **2** (273 mg, 1.0 mmol), 1-bromo-4-vinylbenzene (183 mg, 1.0 mmol), P(*o*-tol)_3_ (61 mg, 20 mol%), triethylamine (0.40 mL, 2.9 mmol) in degassed CH_3_CN (15 mL) under argon was added Pd(OAc)_2_ (23 mg, 10 mol%) quickly. The resulting mixture was stirred at 100 ^°^C for 12 h. The mixture was then diluted with water (20 mL) and the aqueous phase was extracted with ethyl acetate (15 mL × 3). The combined organic extracts were dried over anhydrous Na_2_SO_4_, filtered and evaporated under reduced pressure. Silica flash column chromatography (ethyl acetate/hexanes = 1:30) gave a pale green solid (234 mg, 71%) as the product (**3**). Spectral data of **3** is consistent with those reported in the literature.^(Lemercier et al., 2006)^ To a stirred solution of **3** (165 mg, 0.50 mmol), 4-vinlypyridine (105 mg, 1.0 mmol), P(*o*-tol)_3_ (30 mg, 20 mol%) and triethylamine (0.20 mL, 1.5 mmol) in degassed CH_3_CN (10 mL) under argon was added Pd(OAc)_2_ (11 mg, 10%) quickly. The resulting mixture was stirred at 100 ^°^C for 12 h. The mixture was then diluted with water (20 mL) and the aqueous phase was extracted with ethyl acetate (15 mL × 3). The combined organic extracts were dried over anhydrous Na_2_SO_4_, filtered and evaporated under reduced pressure. Silica gel flash column chromatography (ethyl acetate/hexanes = 3:1) of the residue gave a pale-yellow solid (128 mg, 72%) as the product. PC4: mp = 225-226 ^°^C. ^1^H NMR (400 MHz, CDCl_3_) δ 8.54 (d, *J* = 5.7 Hz, 2H), 7.54 – 7.44 (m, 4H), 7.42 – 7.38 (m, 2H), 7.36 (d, *J* = 6.0 Hz, 2H), 7.29 (d, *J* = 16.3 Hz, 1H), 7.09 (d, *J* = 16.2 Hz, 1H), 6.99 (d, *J* = 16.3 Hz, 1H), 6.88 (d, *J* = 16.2 Hz, 1H), 6.67 (d, *J* = 8.9 Hz, 2H), 3.39 (q, *J* = 7.0 Hz, 4H), 1.19 (t, *J* = 7.0 Hz, 6H). ^13^C NMR (100 MHz, CDCl_3_) δ 150.1, 147.6, 144.9, 139.1, 134.2, 133.1, 129.7, 129.0, 128.6, 128.0, 127.4, 126.3, 124.9, 124.4, 122.9, 120.8, 111.7, 44.4, 12.7. HRMS (+ESI) m/z calcd. for C_25_H_26_N_2_ (M+H)^+^ 355.2169, found 355.2167.

To synthesize PC5, to a stirred solution of 4-vinylpyridine (210 mg, 2.0 mmol), 4-iodophenol (220 mg, 1.0 mmol), P(*o*-tol)_3_ (61 mg, 20 mol%) and triethylamine (0.40 mL, 2.9 mmol) in degassed CH_3_CN (15 mL) under argon was added Pd(OAc)_2_ (23 mg, 10 mol%) quickly. The resulting mixture was stirred at 100 °C for 12 h. The mixture was then diluted with water (20 mL). Upon addition of 5% HCl leads to partial precipitation of the product. The aqueous phase was extracted with ethyl acetate (15 mL × 3). The combined organic extracts were dried over anhydrous Na_2_SO_4_, filtered and evaporated under reduced pressure. Silica gel flash column chromatography (ethyl acetate/hexanes = 3:1) of the residue gave an off-white solid (130 mg, 66%) as the product. **PC5**: mp = 281-282. ^1^H NMR (400 MHz, DMSO-*d*_6_) δ 9.83 (s, 1H), 8.55 (d, *J* = 4.5 Hz, 2H), 7.67 – 7.42 (m, 5H), 7.07 (d, *J* = 16.4 Hz, 1H), 6.85 (d, *J* = 9.3 Hz, 2H). ^13^C NMR (100 MHz, DMSO-*d*_6_) δ 158.2, 149.8, 144.8, 133.1, 128.7, 127.2, 122.4, 120.5, 115.7. HRMS (+ESI) m/z calcd. for C_13_H_11_NO (M+H)^+^ 198.0913, found 198.0913.

To synthesize PC6, to a stirred solution of 4-vinylpyridine (631 mg, 6.0 mmol), 1-ethoxy-4-iodobezene (1.24 g, 5.0 mmol), P(*o*-tol)_3_ (305 mg, 20 mol%) and triethylamine (2.0 mL, 15 mmol) in degassed CH_3_CN (15 mL) under argon was added Pd(OAc)_2_ (112 mg, 10 mol%) quickly. The resulting mixture was stirred at 100 °C for 12 h. The mixture was then diluted with water (20 mL) and the aqueous phase was extracted with ethyl acetate (15 mL × 3). The combined organic extracts were dried over anhydrous Na_2_SO_4_, filtered and evaporated under reduced pressure. Silica gel flash column chromatography (ethyl acetate/hexanes = 3:1) of the residue gave an off-white solid (958 mg, 85%) as the product. PC6: mp = 146-147 °C. ^1^H NMR (400 MHz, CDCl_3_) δ 8.55 (dd, *J* = 4.6, 1.5 Hz, 2H), 7.51 – 7.42 (m, 2H), 7.33 (dd, *J* = 4.6, 1.5 Hz, 2H), 7.27 (t, *J* = 8.1 Hz, 1H), 6.97 – 6.81 (m, 3H), 4.07 (q, *J* = 7.0 Hz, 2H), 1.44 (t, *J* = 7.0 Hz, 3H). ^13^C NMR (100 MHz, CDCl_3_) δ 159.6, 150.1, 145.0, 132.8, 128.7, 128.4, 123.6, 120.6, 114.8, 63.6, 14.8. HRMS (+ESI) m/z calcd. for C_15_H_15_NO (M+H)^+^ 226.1226, found 226.1226.

To synthesize PC7, to a stirred solution of 4-iodophenol (1.09 g, 4.93 mmol) and triethylamine (749 mg, 7.40 mmol) in CH_2_Cl_2_ (25 mL) was added acetyl chloride (465 mg, 5.92 mmol) at rt. The resulting mixture was stirred at 0 °C for 20 min and then rt for 2 h. The solution was then diluted with water (20 mL) and the aqueous phase was extracted with ethyl acetate (15 mL × 3). The combined extracts were dried over anhydrous Na_2_SO_4_, filtered and evaporated under reduced pressure. The resulting pale brown oil (1.15 g, 89 %) was obtained as the product (**4**) and was used for the next step without any further manipulation. Spectral data of **4** is consistent with those reported in the literature.^(Flaherty et al., 2010)^ To a stirred solution of 4-vinylpyridine (210 mg, 2.0 mmol), 4 (240 mg, 1.0 mmol), P(*o*-tol)_3_ (61 mg, 20 mol%) and triethylamine (0.40 mL, 2.9 mmol) in degassed CH_3_CN (15 mL) under argon was added Pd(OAc)_2_ (23 mg, 10 mol%) quickly. The resulting mixture was stirred at 100 °C for 6 h. The mixture was then diluted with water (30 mL) and the aqueous phase was extracted with ethyl acetate (15 mL × 3). The combined organic extracts were dried over anhydrous Na_2_SO_4_, filtered and evaporated under reduced pressure. Silica gel flash column chromatography (ethyl acetate/hexanes = 3:1) of the residue gave a white solid (103 mg, 43%) as the product. PC7: mp = 152-153 °C. ^1^H NMR (400 MHz, CDCl_3_) δ 8.58 (d, *J* = 5.9 Hz, 2H), 7.63 – 7.48 (m, 2H), 7.36 (dd, *J* = 4.7, 1.4 Hz, 2H), 7.28 (d, *J* = 16.3 Hz, 1H), 7.15 – 7.09 (m, 2H), 6.97 (d, *J* = 16.3 Hz, 1H), 2.32 (s, 3H). ^13^C NMR (100 MHz, CDCl_3_) δ 169.4, 150.9, 150.2, 144.5, 133.9, 132.2, 128.1, 126.2, 122.1, 120.9, 21.2. HRMS (+ESI) m/z calcd. for C_15_H_13_NO_2_ (M+H)^+^ 240.1019, found 240.1018.

To synthesize PC8, to a stirred solution of 5-bromo-2-hydroxy-benzonitrile (0.60 g, 30 mmol) in DMF (7 mL) was added ethyl bromide (0.37 mL, 5.0 mmol), and K_2_CO_3_ (1.0 g, 9.4 mmol) at rt. The resulting mixture was stirred at 70 °C for 6 h. The mixture was then diluted with water (20 mL), and the aqueous phase was extracted with ethyl acetate (15 mL × 3). The combined organic extracts were washed with water (15 mL × 3), dried over anhydrous Na_2_SO_4_, and evaporated under reduced pressure. A white solid was obtained as the product (**12**). To a stirred solution of the crude product (**12**), 4-vinylpyridine (315 mg, 3.0 mmol), P(*o*-tol)_3_ (183 mg, 20 mol%) and triethylamine (1.2 mL, 8.7 mmol) in degassed CH_3_CN (30 mL) under argon was added Pd(OAc)_2_ (67mg, 10 mol%) quickly. The resulting mixture was stirred at 100 °C for 5 h. The mixture was then diluted with water (30 mL) and the aqueous phase was extracted with ethyl acetate (15 mL × 3). The combined organic extracts were dried over anhydrous Na_2_SO_4_, filtered and evaporated under reduced pressure. Silica gel flash column chromatography (ethyl acetate/hexanes = 3:1) of the residue gave a pale-yellow solid (433 mg, 58%) as the product. **PC8**: mp = 114-115 °C. ^1^H NMR (400 MHz, CDCl_3_) δ 8.59 (d, *J* = 5.9 Hz, 2H), 7.80 – 7.60 (m, 2H), 7.35 (d, *J* = 5.9 Hz, 2H), 7.19 (d, *J* = 16.3 Hz, 1H), 6.99 (d, *J* = 8.8 Hz, 1H), 6.91 (d, *J* = 16.3 Hz, 1H), 4.19 (q, *J* = 7.0 Hz, 2H), 1.50 (t, *J* = 7.0 Hz, 3H). ^13^C NMR (101 MHz, CDCl_3_) δ 160.4, 150.0, 143.9, 132.5, 131.8, 130.2, 128.9, 125.7, 120.6, 115.9, 112.3, 102.4, 64.8, 14.3. HRMS (+ESI) m/z calcd. for C_16_H_14_N_2_O (M+H)^+^ 251.1179, found 251.1178.

To synthesize PC9, to a stirred solution of 1-ethoxy-4-iodobezene (1.24g, 5.0 mmol) and 3,3-diethoxyprop-1-ene (1.03 g, 7.9 mmol), P(*o*-tol) (305 mg, 20 mol%), Cs_2_CO_3_ (2.28g, 7.0 mmol) and KCl (370 mg, 5mmol) in DMF (30 mL) under argon was added Pd(OAc)_2_ (115 mg, 10 mol%) quickly. The resulting mixture was stirred at 90 °C for 5 h and then treated with 5% HCl (10 mL) and stirred at rt for 10 min. The mixture was then diluted with water (20 mL) and the aqueous phase was extracted with ethyl acetate (15 mL × 3). The combined organic extracts were dried over anhydrous Na_2_SO_4_, filtered and evaporated under reduced pressure. Silica gel flash column chromatography (ethyl acetate/hexanes = 1:5) gave a pale-yellow solid (443 mg, 50 %) as the product (**6**). Spectral data of **6** is consistent with those reported in the literature.^(Lator et al., 2018)^ To a stirred solution of **6** (88 mg, 0.50 mmol), 4-methylpyridine (93 mg, 1.0 mmol) in Ac_2_O (2 cmL) was added NaOAc (272 mg, 2.0 mmol) at rt. The resulting mixture was heated under reflux for 21 h. Then the mixture was cooled to rt and diluted with CH_2_Cl_2_, washed with H_2_O, 5% HCl, H_2_O and saturated aqueous NaHCO_3_. The aqueous phase was extracted with ethyl acetate (15 mL × 3). The combined organic extracts were dried over Na_2_SO_4_, filtered and evaporated under reduced pressure. Silica gel flash column chromatography (ethyl acetate/hexanes = 3:1) gave a pale-yellow solid (47 mg, 37%) as the product. **PC9**: mp = 132-133 °C. ^1^H NMR (400 MHz, CDCl_3_) δ 8.58 (d, *J* = 5.9 Hz, 2H), 7.44 (d, *J* = 8.7 Hz, 2H), 7.36 – 7.29 (m, 2H), 7.17 (dd, *J* = 15.5, 10.2 Hz, 1H), 6.99 – 6.74 (m, 4H), 6.57 (d, *J* = 15.5 Hz, 1H), 4.11 (d, *J* = 7.0 Hz, 2H), 1.48 (t, *J* = 7.0 Hz, 3H). ^13^C NMR (101 MHz, CDCl_3_) δ 159.0, 149.8, 144.7, 135.4, 133.8, 129.2, 128.2, 127.9, 125.8, 120.3, 114.6, 63.3, 14.6.HRMS (+ESI) m/z calcd. for C_17_H_18_NO (M+H)^+^ 252.1383, found 252.1384.

### Preparation and quantification of the enzymes

A truncated form of SARM1, SARM1-dN, was prepared as described.(Zhao et al., 2019) In brief, the recombinant protein was expressed in HEK293T cells and released by 100 μM digitonin in PBS with protease inhibitor cocktail (Roche). The cell lysate prepared with the same method from HEK293T was used as the negative control.

To quantify the protein concentration, SARM1-dN was pulled down by BC2 nanobody(Bruce et al., 2017) conjugated beads which were prepared by conjugating BC2 nanobody to NHS-beads (GE Healthcare). The purified SARM1-dN, named as SARM1-IP, together with the certain amounts of standard protein BSA, was applied to SDS-PAGE, which was stained by Coomassie blue. The protein contents of SARM1-dN was calculated with a standard curve of BSA after intensity scanning by Image J.

Recombinant CD38 and N. crassa NADase were prepared as described previously.(Graeff et al., 1994, Munshi et al., 1997)

### *In vitro* fluorescence assays

To analyze the activity of SARM1 with PCs in vitro, reactions were started by incubating the enzyme with the reaction mixture, 50 μM PC, 100 μM NAD and 100 μM NMN in PBS. The absorbance and fluorescence were measured in a quartz cuvette or black 96-well plates (Corning), respectively, with an Infinite M200 PRO microplate reader (Tecan). For the assays with εNAD, NHD or NGD as the substrate, 100 μM the compounds were incubated with the enzymes and fluorescence were monitored at λ_ex_ = 300 nm, λ_em_ = 410 nm.

### HPLC analysis of the base-exchange reaction of PC6 catalyzed by SARM1

The reactions were prepared by mixing SARM1-IP (SARM1 binding on BC2-beads, around 2.5 μg/ml) with 100 μM NAD, 50 μM PC6, 100 μM NMN and 0.1 mg/ml BSA in PBS and incubated for 60 min at 37 °C. SARM1-IP was removed by centrifugation at 4,500 rpm for 1 min. The cleaned mixtures were applied to a C-18 reverse phase column equipped on an HPLC (Agilent 1260) with a gradient of 0.1 M KH_2_PO_4_ (pH 6.0) and 0.1 M KH_2_PO_4_ (pH 6.0) with MeOH (7:3) to elute NMN, cAPPR, ADPR, NAD and a gradient of ACN from 30% to 70% to elute PAD6 and PC6. The fractions of PAD6 were collected and lyophilized for absorption spectra and fluorescence spectra scanning.

To analyze the PAD6 in cells, the metabolites were extracted from the cells treated with 50 μM PC6 by 0.6 M perchloric acid, followed by neutralization with Chloroform: Tri-n-octylamine (3:1). The extracts were applied to a C-18 column and eluted with water and acetonitrile by 2% acetonitrile for 8 min then 30% acetonitrile for 8 min.

### Confocal imaging of PAD6 in living cells

HEK293 cells, overexpressing wildtype or the enzymatically dead form (E642A) of SARM1 or HEK293T Knocking out NAMAT1 were constructed as before.(Zhao et al., 2019) Cells, grown on 0.05 mg/ml poly-L-lysine coated Chambered coverglass (Thermo fisher, #155411) overnight were treated with 50 μM of PC6 in the presence or absence of 100 μM CZ-48 for 8 h (SARM1-OE cells) and 200 μM CZ-48 for 48 h (original cell lines), respectively. To demonstrate the edges of the cells, they were stained with 50 μg/ml Concanavalin A, Alexa Fluor™ 647 Conjugate (Thermo fisher) at 4°C for 10 min before imaging. The fluorescence signals (Ex/Em: 405/520 nm for PAD6; Ex/Em: 561/590 for ConA) were captured under a confocal microscope (Nikon A1).

### Analysis of PAD6 signals in living cells by flow cytometry

HEK293 cells carrying an inducible expression cassette of SARM1 were constructed as previous describe.(Zhao et al., 2019) The cells were treated with 50 μM PC6, 100 μM CZ-48 or 0.5 mg/mL Dox for 4, 8, 12 and 16 h. The cells were trypsinized and the fluorescence of PAD6 (Ex/Em: 405/525 nm) was analyzed by flow cytometry (CytoFlex, Beckman).

### DRG culture and imaging

Mouse DRG culture was performed as described.(Sasaki et al., 2016) Briefly, DRGs were dissected from the embryos at Day 12.5 to 14.5 (E12.5-E14.5) and digested by 0.05% Trypsin solution containing 0.02% EDTA (Gibco). The dispersed cells were seeded in Neurobasal™ Plus Medium supplemented with 2% B27 plus, 1 mM GlutaMax, 1% penicillin/streptavidin solution and 37.5 ng/ml NGF on the Chambered coverglass pre-coated with (0.1 mg/ml) poly-L-Lysine, (0.02 mg/ml) laminin and 5% FBS. Every 3 day, 50 % of the culture media was replaced by fresh media with the addition of 5 μM 5-fluoro-2’-deoxyuridine and 5 μM uridine.

On Div6 the neurons were infected with lentivirus. Three days later, the cells were treated with 50 μM PC6 in the absence or presence of 200 μM CZ-48 or 50 nM vincristine. The fluorescence images (Ex/Em: 405/520 nm for PAD6; Ex/Em: 561/590 for TdTomato) were captured under a confocal microscope (Nikon A1) with a 60x object. The mean fluorescence intensity was quantified by NIS-Elements AR analysis (Nikon). Axon degeneration was quantified base on axon morphology using ImageJ. The TdTomato fluorescence images were binarized and measured the total axon area (size = 20-infinity pixels) and the degenerated axon (size = 20-4,000,000 pixels) with particle analyzer module of ImageJ. Axon degeneration index was calculated as the ratio of the degeneration axon over total axon area.

### Lentivirus preparation and infection of DRG neurons

pLKO.1-shRNA-mSARM1 plasmids were constructed as described previously.(Zhao et al., 2019) Briefly, shRNA targeting mSARM1 (5’-CCGGCTGGTTTCTTACTCTACGAATCTCGAGATTCGTAGAGTAAGAAACCAGTTTTTG-3’) or the scrambled shRNA (5’-CCGGCCTAAGGTTAAGTCGCCCTCGCTCGAGCGAGGGCGACTTAACCTTAGGTTTTTG-3’) were inserted to pLKO.1-puro (Addgene, #8453) with EcoRI and AgeI, followed by replacement of the puromycin resistance gene with a fluorescent protein, TdTomato (GenBank: LC311026.1) with KpnI/BamHI. The lentiviral particles were prepared by transfecting HEK293T cells with the corresponding lentivectors, pMD2.G and psPAX26,(Liu et al., 2017) followed by concentration of the virus with Lenti-Concentin Virus Precipitation Solution (ExCell Bio), which was resuspended in Neurobasal™ Plus Medium. Determine the virus titer by series infection of HEK293T cells. For infection of DRG neurons, added the same MOI of virus to infect the cells on Div 6 and carried out further experiment 72 hours after infection.

### Imaging and quantification of AxD after axotomy and vincristine treatment

For axotomy, one DRG was seeding into a 24-well plate, and 5 μM 5-fluoro-2’-deoxyuridine and 5 μM uridine was added the other day. On div5, axons were pre-incubated with drugs for 0.5 h and severed near the soma with a 3 mm flat blade under microscope guidance removing the cell bodies. For vincristine treatment, DRGs was digested with 0.05% Trypsin and seeding into 24-well plate. DRGs on Div9-13 were incubated 50 nM vincristine in the presence or absence of the drug.

9-12 brightfield images per treatment of the axon were acquired with a 20x object at the indicated time points using invert optical microscope (Olympus). Axon degeneration was quantified base on axon morphology using ImageJ. For each treatment, 60 random grid-squares with 147×147 pixels were cropped, binarized and measured the total axon area (size = 16-infinity pixels) and the degenerated axon (size = 16-10,000 pixels) with particle analyzer module of ImageJ. Axon degeneration index was calculated as the ratio of the degeneration axon over total axon area.

### Measuring cADPR level of DRG treating with vincristine

DRG neurons were treated with 50 nM Vincristine or 200 μM CZ-48 for 0, 12, 24, 48 h on Div6. After incubation, DRG was washed with cold PBS and lysed with 0.6 M perchloric acid. cADPR was extracted and analyzed as described previously.(Graeff et al., 2002)

### Q-RT-PCR

After infection of virus for 48 h, total RNAs were extracted from DRG neurons with RNA extraction kit (OMEGA) and transcribed with Transcript II One-step gDNA Removal and cDNA synthesis Supermix (Sangon Biotech). The mRNA level of SARM1 relative to GAPDH was quantified with by Q-RT-PCR using TransStart Tip Green qPCR SuperMix (TransGen Biotech) on CFX Connect Real-Time PCR Detection System (Bio-Rad). The following primer pairs were used: SARM1 sense, 5’-CTTTCTCCAAGGAGGACGAGC-3’, antisense, 5’-CTTGTGTCACTGGCATCCACC-3’; GAPDH sense, 5’-TGGCCTTCCGTGTTCCTAC-3’, antisense, 5’-GAGTTGCTGTTGAAGTCGCA-3’.

### PC6 assay

For high-throughput screening, inhibition of 1.5 μg/ml SARM1-dN in vitro determined by 20 μM PC6 with 50 μM of 2046 compound from a drug approval library (Targetmol, L1000) in the presence of 50 μM NAD and 50 μM NMN. Thirty-four compounds with high fluorescence overlapped with PC6 assay were removed.

For IC_50_ measurement, 0.4 μg/ml SARM1-dN was pre-incubated with dose of compounds *in vitro* for 10 min, and started the reaction by adding 50 μM NAD, 50 μM NMN and 50 μM DA4. Calculation of IC_50_ by plotting the initial rate to dose of compounds.

### HPLC analyze SARM1 NADase acitivy

1 μg/ml of SARM1-dN was pre-incubated with compounds for 15 min at room temperature, and started the reaction by adding 100 μM NAD and 100 μM NMN. The reactions were stopped by removing the enzyme with MultiScreen® Filter Plates (Millipore) after 0, 15 and 30 min incubation at 37 °C and analyzed by a C-18 column (Aligent, 20 RBAX SB-C18) with a gradient of 0.1 M KH_2_PO_4_ (pH 6.0) and 0.1 M KH_2_PO_4_ (pH 6.0) with MeOH (7:3) to elute NMN, cAPPR, ADPR, NAD, Nam. The amount of ADPR were used to calculate the initial rate. IC_50_ of NADase activity was calculated by plotting dose of compounds to the initial rate.

### HPLC to analysis nisoldipine and its derivatives

10 mM NSDP was prepared freshly in DMSO and treated with UV at 254 nm for 30 min. 2.5 nmole NSDP-UV and standards were applied to a C-18 reverse phase column (ZORBAX SB-C18) equipped on a HLPC (Aligent 1260) and eluted with 50% of 0.1%TFA and 50% of 0.1%TFA with 99% ACN. 0.3 μmole of the product after UV treatment was collected and purified by HPLC described above. The inhibitory activity of these fractions was determined by PC6 assay after neutralized with 100 mM Tris (pH7.5), and the main peak was characterized by HRMS (thermo, Q Exactive Focus).

### Measure the inhibitory activity of dHNN *in vitro* and *in cellulo*

To determine whether dHNN inhibits activation or enzymatic activity of SARM1 *in vitro*, SARM1-dN, the autoinhibited form, and SAM-TIR, the constitutively active form, were pre-incubated with different concentrations of dHNN at RT for 10 min, after which the activity was measured with PC6 assay and the inhibition rate was calculated.

To test the same effect *in cellulo*, HEK293 cells overexpressing inducible SARM1 (iSARM1) or SAM-TIR (iSAM-TIR) were pre-incubated with 20 μM dHNN for 1.5 h and then treated with 100 μM CZ-48 or 0.5 μg/ml doxycycline for the indicated time. Control cells were similarly induced but pre-incubated with vehicle DMSO. Cellular cADPR of the sample cells was measured and divided by that of the corresponding control cells to determine the percent reduction of cADPR content.

### Modification analysis of SARM1 by dHNN

dtSARM1-dN that had the N-terminal targeting signal removed, was tagged with strep tag II and flag for purification, constructed into pENTR1A-GFP-N2 and cloned into Plenti-CMV-puro-Dest (Invitrogene) by LR clonase II enzyme according to the manufacturer’s instructions. HEK293F cells were infected by lentivirus carrying dtSARM1-dN produced by HEK293T cells with lipo2000 transfection and selected with 1 μg/ml puromycin. The cells were harvested by PBS and the pellet were stocked at −80°C before using. Protein extracted with 200 μM digitoinin were performed immunoprecipitated with StrepTactin beads overnight. The beads were washed with buffer W (pH7.5) containing 100 mM Tris, 150 mM NaCl and 1 mM EDTA for four times and eluted on ice with 2 mM biotin in buffer W. 0.17 mg/ml dtSARM1-dN pre-incubated with 50 μM dHNN at RT for 30 min were boiled in SDS loading buffer and applied to SDS-PAGE gel. The gel was stained by simplyBlue^TM^ SafeStain (Thermo fisher) for 1 h and destained by water for 2 h with water changing for three times. dtSARM1-dN were dissected form the gel, dehydrated with 100% ACN, and performed alkylation by incubated with 10 mM DTT at 55°C for 30 min and 22.5 mM IAA for another 30 min in dark. Then the gel were destained with 50% ACN in 25 mM (NH_4_)_2_SO_4_, dehydrated with ACN and protein were digested in gel with Trypsin at 37°C overnight. Peptide were extracted from the gel with 5% FA and 50% ACN in water, then lyophilized and resuspended with 0.1% FA in water for mass spectrometry identification (Thermo, Q Exactive HF-X). The modification of dHNN on cysteine were analyzed by protein discoverer (Thermo fisher).

### Cysteine mutants

The mutants of cysteine to alanine in dtSARM1-dN, described above, were cloned by overlapping PCR with the primer below by PrimeStar HS polymerase with high GC buffer. The genes were constructed into pCDH-EF1-MCS-IRES-neo by Xba I and Not I, and transfected into HEK293 cells by lipofectamine 2000 or Polyethylenimine according to the manufacturer’s instructions. Proteins were extracted 48-72 h after transfection and determined the IC_50_ with dHNN by PC6 assay *in vitro*.

C117A-F: 5’-GTAGCCCAGGGTCTGGCC GACGCCATCCGC-3’

C117A-R: 5’-GCGGATGGCGTCGGCCAGACCCTGGGCTAC-3’

C199A-F: 5’-CATTCGGAGGAGACAGCC CAGAGGCTGGTG-3’

C199A-R: 5’-CACCAGCCTCTGGGCTGTCTCCTCCGAATG-3’

C215A-F: 5’-GCGGTGCTGTATTGGGCACGCCGCACGGAC-3’

C215A-R: 5’-GTCCGTGCGGCGTGCCCAATACAGCACCGC-3’

C226A-F: 5’-GCGCTGCTGCGCCACGCAGCGCTGGCGCTG-3’

C226A-R: 5’-CAGCGCCAGCGCTGCGTGGCGCAGCAGCGC-3’

C233A-F: 5’-CTGGCGCTGGGCAACGCAGCGCTGCACGGG-3’

C233A-R: 5’-CCCGTGCAGCGCTGCGTTGCCCAGCGCCAG-3’

C271A-F: 5’-CTTCGGCTGCACGCCGCACTCGCAGTAGCG-3’

C271A-R: 5’-CGCTACTGCGAGTGCGGCGTGCAGCCGAAG-3’

C311A-F: 5’-GGCCGCTTCGCCCGCGCC CTGGTGGACGCC-3’

C311A-R: 5’-GGCGTCCACCAGGGCGCGGGCGAAGCGGCC-3’

C343A-F: 5’-CGCTTGGAGGCGCAGGCAATCGGGGCTTTC-3’

C343A-R: 5’-GAAAGCCCCGATTGCCTGCGCCTCCAAGCG-3’

C350A-F: 5’-GGGGCTTTCTACCTCGCAGCCGAGGCTGCC-3’

C350A-R: 5’-GGCAGCCTCGGCTGCGAGGTAGAAAGCCCC-3’

C430A-F: 5’-GGTTTCTCCAAGTACGCAGAGAGCTTCCGG-3’

C430A-R: 5’-CCGGAAGCTCTCTGCGTACTTGGAGAAACC-3’

C482A-F: 5’-GCCAACTATTCTACGGCC GACCGCAGCAAC-3’

C482A-R: 5’-GTTGCTGCGGTCGGCCGTAGAATAGTTGGC-3’

C508A-F: 5’-TACGGCCTGGTCAGCGCAGGCCTGGACCGC-3’

C508A-R: 5’-GCGGTCCAGGCCTGCGCTGACCAGGCCGTA-3’

C527A-F: 5’-CAGCTGCTGGAAGACGCAGGCATCCACCTG-3’

C527A-R: 5’-CAGGTGGATGCCTGCGTCTTCCAGCAGCTG-3’

C552A-F: 5’-CACTCCCCGCTGCCCGCAACTGGTGGCAAAC-3’

C552A-R: 5’-GTTTGCCACCAGTTGCGGGCAGCGGGGAGTG-3’

C629A-F: 5’-GGAGCACTGGACAAGGCAATGCAAGACCAT-3’

C629A-R: 5’-ATGGTCTTGCATTGCCTTGTCCAGTGCTCC-3’

C635A-F: 5’-ATGCAAGACCATGACGCAAAGGATTGGGTG-3’

C635A-R: 5’-CACCCAATCCTTTGCGTCATGGTCTTGCAT-3’

C649A-F: 5’-GTGACTGCTTTAAGCGCC GGCAAGAACATT-3’

C649A-R: 5’-AATGTTCTTGCCGGCGCTTAAAGCAGTCAC-3’

dtSARM1-dN-F: 5’-CAGTCTAGAATGGACTACAAGGATGACGATG-3’

dtSARM1-dN-R: 5’-ATAGCGGCCGCTTAGGTTGGACCCA-3’

### Western blots

Cells was harvested and lysed with RIPA buffer (50 mM Tris-HCl, 150 mM NaCl, 1 mM EDTA and 0.05% triton, pH 7.4). Each sample was loaded onto 10-12% SDS-PAGE gels and transferred to PVDF membranes after electrophoresis. The membranes were blocked with 5% Milk and blotted with anti-SARM1 (produced as described previously) and anti-Tubulin antibody (TransGen Biotech) to control the loading.

### CryoEM sample preparation, data collection and processing

SARM1-dN tagged with strep-tag II and flag-tag was immunoprecipitated with StrepTactin resin (GE healthcare), washed with buffer W (100 mM Tris-HCl pH8.0, 150 mM NaCl and 1 mM EDTA) for four times and eluted with 2 mM biotin in buffer W. The eluent was concentrated to 3 mg/ml and pre-incubated with 50 μM dHNN at RT for 10 min.

SARM1-dHNN protein was applied to glow-discharged gold grid, blotted in FEI Vitrobot Mark IV (Thermo Fisher Scientific) before frozen by liquid ethane and stored in liquid nitrogen. The sample without inhibitor was examined at the Cryo-EM center of Chinese University Hong Kong (Shenzhen) on a 300kV Titan Krios (Thermo Fisher Scientific) equipped with Gatan K3 direct electron detector under magnification of 105,000x, with the corresponding pixel size of 0.85Å. The dose rate was set to 17.6 e/pix/s and exposure time was set to 2.5s to obtain 50 frames, which led to an accumulated dose of 61 electrons per Å2. The total dataset consists of 2,692 raw movies with a defocus value range of −1.0 to −2.0 μm. Motion correction and CTF parameter estimation were performed with cryoSPARC (Punjani et al., 2017). 2,012,198 particles were autopicked. After several rounds of 2D classification, 712,139 particles were selected for generation of the final 2D average results.

The dHNN-treated sample was examined at the Cryo-EM center of Southern University of Science and Technology on a Titan Krios G3 (Thermo Fisher Scientific) with Gatan K2 summit detector with a nominal magnification of 130,000X and corresponding pixel size of 1.076 Å. A total accumulative dose of 50 e^−^/Å2 was set for each exposure and split into 39 frames during data acquisition. The defocus range was set between −0.8 to −2.0 μm. In total, 2,890 images were collected. Motion correction and CTF parameter estimation were performed with MotionCor2 and CTFFind4 built within Relion 3.1(Fernandez-Leiro et al., 2017). After CTF estimation, images with thick ice, obvious shift or cleft were removed, which left 2673 images for further processing. 2,655,835 particles were autopicked from these images. After several rounds of 2D classification, 700,472 particles were selected and exported for generation of the final 2D average results with CryoSparc and 3D refinement with CisTEM beta-1.0.0 (Grant et al., 2018). The particle stack was subject for 10 rounds of 3D auto-refinement among 6 classes using 6WPK as initial model. Four classes with higher estimated resolution were selected and combined for 20 more rounds of 3D manual global refinement and one class with the highest occupation (62.5%) and best resolution was chosen for several more rounds of 2D and 3D classification with Relion 3.1 and CisTEM beta-1.0.0. The resolution for the final map was around 2.4 Å..

The previously reported structures of the SARM1 SAM domain (PDB: 6O0S) and TIR domain (PDB: 6O0Q) were used as model templates during initial model building. The initial model of ARM domain was built de novo in Coot (Emsley et al., 2010). The three domains of SARM1 were connected in Coot and docked into density maps using Dock in Map module of Phenix 1.16 (Adams et al., 2010) with C8 symmetry and then subjected to multiple rounds of Real-space refinement in Phenix. The dHNN molecule was built and fitted into the density around Cys311 initial model in Coot. The final models were validated with Comprehensive Validation module of Phenix and the refinement statistics were listed in Supplementary file 1. The model and EM map have been deposited in Protein Data Bank with accession codes of PDB ID 7DJT and EMD-30700.

### Data analysis

All experiments contained at least three biological replicates. Data shown in each figure are all means ± SD. The unpaired Student’s t-test was used to determine statistical significance of differences between means (*P<0.05, **P<0.01, ***P<0.001, ****P<0.0001). GraphPad Prism 7 was used for data analysis.

## ACKNOWLEDGMENTS

We would like to thank the Cryo-EM center of Southern University of Science and Technology for Cryo-EM data collection and the HPC-Service Station in Cryo-EM center of Southern University of Science and Technology for data processing. We acknowledge Beijing Artivila Biopharma Co. Ltd for the supports.

